# Mapping axon diameters and conduction velocity in the rat brain – different methods tell different stories of the structure-function relationship

**DOI:** 10.1101/2023.10.20.558833

**Authors:** Christian S. Skoven, Mariam Andersson, Marco Pizzolato, Hartwig R. Siebner, Tim B. Dyrby

## Abstract

The structure-function relationship of nerve fibers describes an empirically determined linear relationship between axon diameter, myelin thickness (i.e., g-ratio), and conduction velocity. We investigated the structure-function relationship with different modalities in axons projecting through the corpus callosum of the rodent brain. We measured transcallosal conduction times using optogenetically evoked local field potentials (LFP) and estimated conduction velocity after measuring the callosal length with diffusion magnetic resonance imaging (dMRI) based tractography. Tractography followed the same projection as the fluorescently labeled axons in the corpus callosum. In the same animal, axon diameters were quantified using transmission electron microscopy (TEM) and dMRI. Axon distributions of TEM indicated a bimodal population, where the larger axons shrink more than the smaller ones, when comparing the modes with cryo-TEM. When applying shrinkage-correction to axon diameters from TEM of dehydrated tissue, they were better aligned with estimates from dMRI obtained in the same animal. Measured LFPs predicted axon diameters that agree with the primary mode of the axon distribution, whereas the large axons estimated by dMRI predicted latencies too short to be measured by LFPs. Different modalities show different degrees of variations, being low between animals, suggesting that the variation is methodologically dominated - not anatomically. Our results show that modalities have different sensitivity profiles to the whole axon diameter distribution. Therefore, caution must be taken when interpreting a method’s prediction of a metric, as it may not represent the full but only a sub-part of the structure-function relationship of an axonal projection.

**Significance Statement:** We acquire both functional and structural metrics, namely conduction velocity, pathway length, axon diameter, and g-ratio in the same cohort of animals. The transcallosal conduction time was obtained from electrophysiological measurements after contralateral optogenetic stimulation in the rat motor cortex. Cryo-fixation of the tissue reveals different shrinkages for different sub-populations in the diameter distribution. Measured latencies correspond to the small axonal subpopulation with diameters extending up to the mode of the distribution obtained with electron microscopy. Diffusion-MRI primarily appears sensitive to the larger axons, obtained with histology, after correcting for diameter weighting and shrinkage. Different modalities can have very different sensitivities to the structure-function relationship of an axonal projection which must be accounted for in the interpretation.

## 1. Introduction

The microstructural morphological properties of the axonal bundles forming the long-distance white matter tracts between functional regions, modulate saltatory signal conduction velocity and in turn the timing of brain function. Rushton et al. [1] put forward a formal theory to describe the structure-relationship between axon diameter, myelin thickness, and conduction velocity [1–4]. The ratio between axon diameter and myelin thickness, referred to as “g-ratio”, is relatively constant in the CNS across different animal species, [5–9]. Assuming that axons have a cylindrical shape and an invariant g-ratio, the conduction velocity is directly proportional to diameter, as is supported by both experiments [2] and simulations [9] [10]. Although other physiological factors also impact conduction velocity [1,2,11,12], it is the axon diameter and myelin that carries the most weight in determining conduction velocity [12].

The distribution of axon diameters varies among brain white matter tracts and depends on which brain regions are connected by the tract [7], as seen in the corpus callosum (CC), which contains most of the inter-hemispheric fibers of most mammalian species [13–15]. Light microscopy (LM) and electron microscopy (EM) of the CC have consistently shown that the axon diameter distributions vary across the midsagittal plane where the largest are related motor- and somatosensory areas [16–19]. By measuring the tract length from injected neuronal tracers or diffusion magnetic resonance imaging (dMRI) based tractography and assuming constant a g-ratio Caminiti et al. predicted in monkeys the conduction latencies between interconnected regions from conduction velocity estimates [19,20]. Good agreement was obtained between the predicted transcallosal latencies and those obtained with electrophysiological recordings - and likewise stimulation experiments in other animal studies [20,21].

*In vivo,* it is possible to obtain an estimate of axon diameter with dMRI, which exploits the restricted diffusion of water molecules to non-invasively probe the geometry of axons [22,23]. The diameter estimation relies on fitting a biophysical model to the measured dMRI signal [22,24]. Horowitz et al. estimated the axon diameter distribution in living humans and correlated the mean diameter index with conduction velocities obtained from electroencephalography [25]. However, the dMRI-based diameters overestimate the values expected from histology [22–24]. The range depends on the MRI hardware and imaging protocol [26–29] which sets an upper and lower bound of measurable axon diameters [26,29] meaning the smallest cannot be measured. Moreover, contrary to a simple arithmetic mean of the diameter distribution as done in 2D histology, the diameter estimate from dMRI is weighted towards large diameters, due to the nature of the scaling of signal attenuation and MRI being a volumetric measure [22,30–33].

Tissue preparation for classical resin-embedding for Electron Microscopy (EM) involves a dehydration step, introducing shrinkage to the tissue between 0-65% [16,23]. Cryo-fixation tissue preparation in comparison introduces minimal shrinkage, approaching realistic measurements of axon diameters in the *in vivo* state [34].

This study aims to explore the structure-function relationship, described by Hursh [2] and others [10], of CNS axons with different measurements of structure and function from the same animals. Our approach is correlating functional or structural measures to their counterparts from other studies. Specifically, we aimed to compare invasive and non-invasive structural imaging modalities (EM, Cryo-EM, and dMRI) to measure the axon and myelin diameters, and to determine their structure-function correlates with electrophysiological recordings.

## 2. Results

### 2.1. Recording the Transcallosal Conduction Time

The latencies from three transcallosal evoked responses i.e., the P1 peak, N1 onset, and N1 peak latencies, as outlined in Fig. 1a, were recorded in 16 animals (Supplementary Table 1). We obtained the recordings under stimulation conditions that produced a reliable N1 peak, which typically included higher stimulation intensity (≥ 1 mW) and/or longer stimulation durations (≥ 1 ms). The mean latency ± mean of *within*-animal standard deviation (± SD) was 5.6 ± 0.3 ms for P1 (N=13), 7.2 ± 0.3 ms for the N1 onset (N=14) and 11.8 ± 0.5 ms for the N1 peak (N=14), respectively, as shown in Fig. 1b. The P1 peak was the earliest transcallosal response that was consistently detectable across most animals (N=13), and was thus used as an indicator of the transcallosal conduction time (TCT). In comparison to the *within*-animal SD of the latencies, the *between*-animal SDs were relatively large, being 1.49, 1.81, and 1.84 ms for the P1 peak, N1 onset, and N1 peak, respectively. Hence, higher variability between animals than within. As the variation between animals might reflect differing electrode depths, we measured the electrode depths on high-resolution structural T2-weighted MR images. The mean (± SD) depth of 1045.23 ± 47.55 µm (N=5; Supplementary Table 2) suggests that depth variation was negligible.

**Figure 1:**
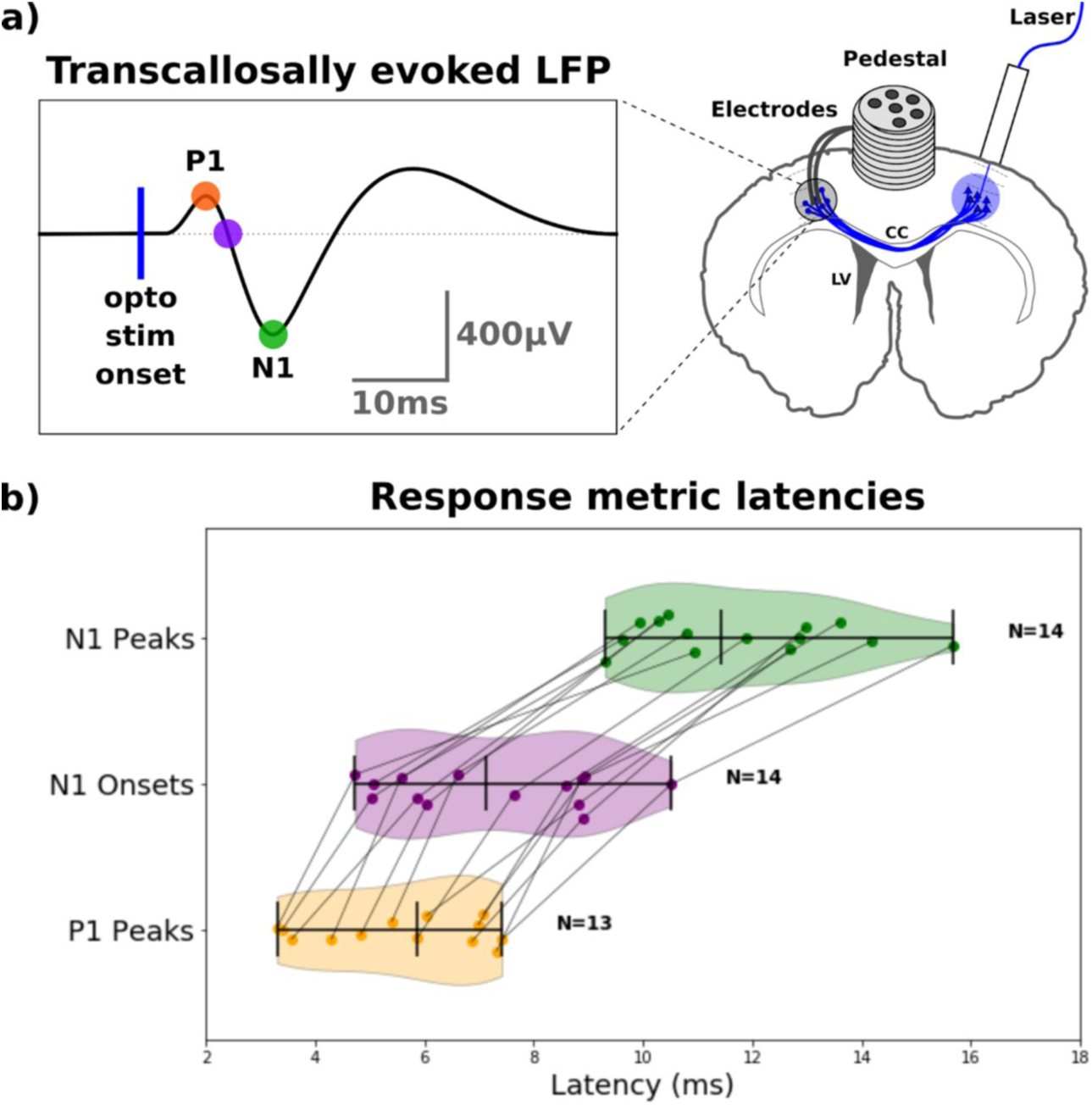
Conduction latencies obtained through optogenetic stimulation and electrophysiology, measured as local field potentials (LFP). a) Optogenetic stimulation of the right M1 and electrophysiological recording in the left M1. Orange dot: P1 peak; Purple dot: N1 onset; Green dot: N1 peak. **b)** Plot of median values of peak detections for all animals [36] in reliable conditions, with a coefficient of variation for the N1 peak being < 1.00. Colors correspond to the insert in subfigure (a). The group means are depicted as the middle vertical line in each of the three metrics, and the outer-most vertical lines represent the minimum and maximum values for each metric.

### 2.2. Mapping the pathway length to estimate the conduction velocity

The viral injection in the right primary motor cortex (M1) targeted excitatory neurons (through the CamKIIα promoter) projecting through the CC to the contralateral M1 (Fig. 2a). Since these optogenetically stimulated neurons (Fig. 1) were fluorescence-labeled (Fig. 2b), their trajectories could be detected through the mid-body of the CC and was present in the superior half of the anterior midbody of CC (Fig. 2b, magnified in the red box insert).

**Figure 2:**
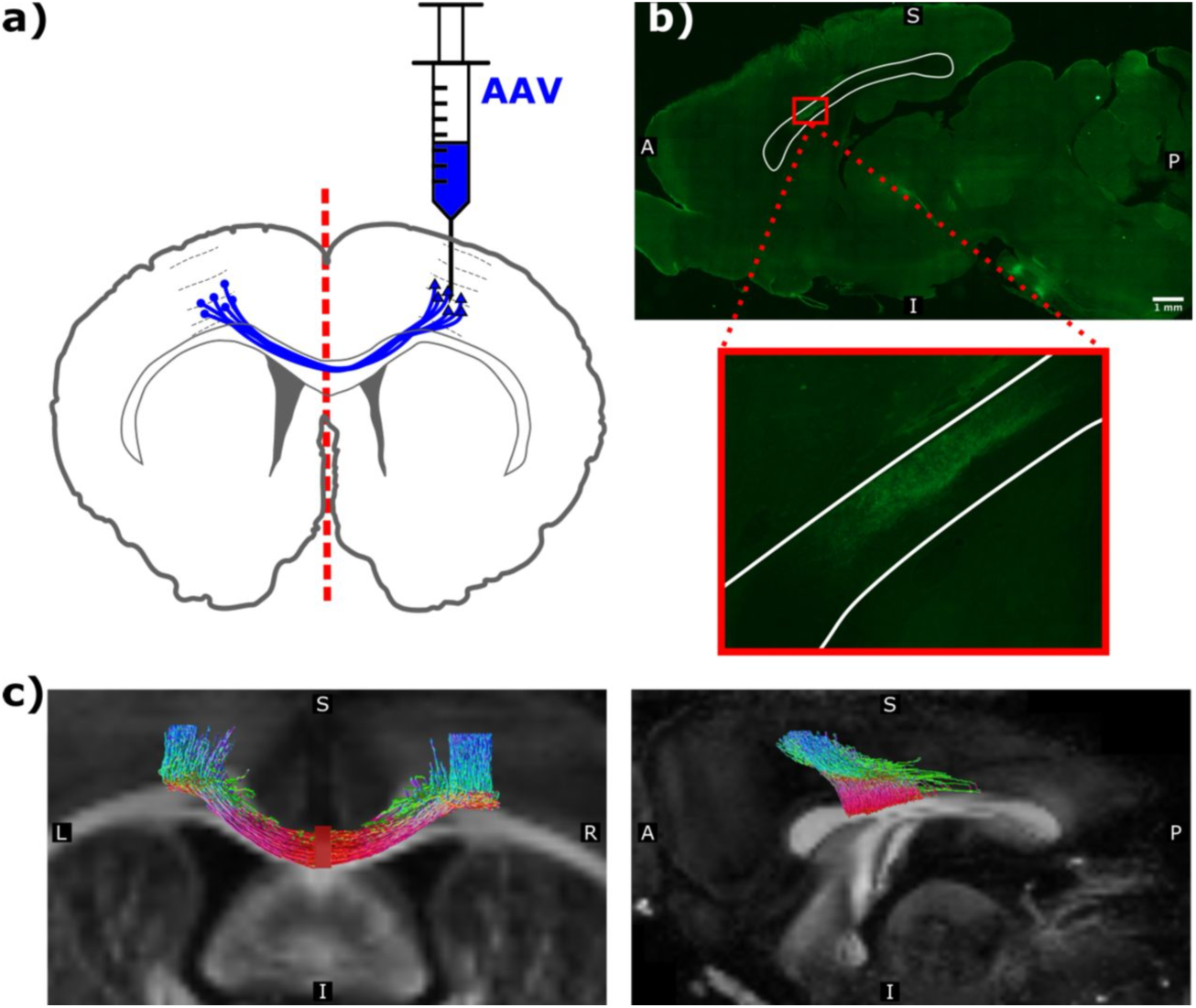
Viral expression and mapped projection pathway of the corpus callosum of rat14.5. a) Illustration of the viral injection in the right M1. **b)** Digitally stitched mid-sagittal image composed of multiple 10x magnification images acquired by fluorescence microscopy traversing the whole slice. Green corresponds to the expression of enhanced yellow fluorescent protein (EYFP) from the virally transfected neurons, highlighting the projection pathway between the primary motor cortices through the CC. The insert figure is an enlarged view of the small red box in the top panel. White lines delineate the contour of the mid-sagittal CC. Scale bar corresponds to 1 mm. **c)** Coronal (left panel) and sagittal (right panel) view of the dMRI scan of rat33.1, overlayed with the tractography streamlines projecting between the seeding region (located in the left M1 and right M1), projecting towards and terminating in the mid-sagittal region (filled red rectangle in the coronal section, left). The color-coding of tractography streamlines indicates their local direction, with red being left/right, green being anterior/posterior, and blue being superior/inferior.

Fig. 2c shows that tractography in the same animal was able to reconstruct the interhemispheric connection between the stimulation (right M1) and the recording sites (left M1). Indeed, a visual inspection of Fig. 2c confirmed that the tractography overlapped with the CC projection region from fluorescence-labeled axons. The tractography-derived connection length across all animals was 11.47 ± 0.47 mm (mean ± SEM; N=8; Supplementary Table 3).

### 2.3. Measuring the axon diameters and g-ratios

Conduction velocities were predicted from structural measurements, using equation (2) in Methods. In a sub-group of the animals (N=4), axon diameter estimates of the same brain were obtained from both *post mortem* dMRI and subsequently axon diameter and g-ratio measures from Epon-TEM images. Some animals were exposed to only either EM (N=1) or dMRI (N=3).

We prepared an Epon-embedded sample from the midsagittal CC, where fluorescently labeled axons projected through (Fig. 2b), for TEM examination (Fig. 3a and 3b). Axon diameters and g-ratios were segmented automatically using AxonDeepSeg [35].

Figures 3c and 3d show the axon segmentation results from a representative case among 79 TEM images from rat14.1, and the corresponding histogram of the axon diameter distribution (N = 853 axons). The gamma distribution of the axon diameter distribution has a mode of 0.48 µm (mean = 0.52 µm; var = 0.02; skew = 0.57), with few axons being larger than 1.0 µm. The group mean (± SD) of the distribution modes was 0.36 ± 0.04 µm (N=5; Supplementary Fig. 2).

**Figure 3:**
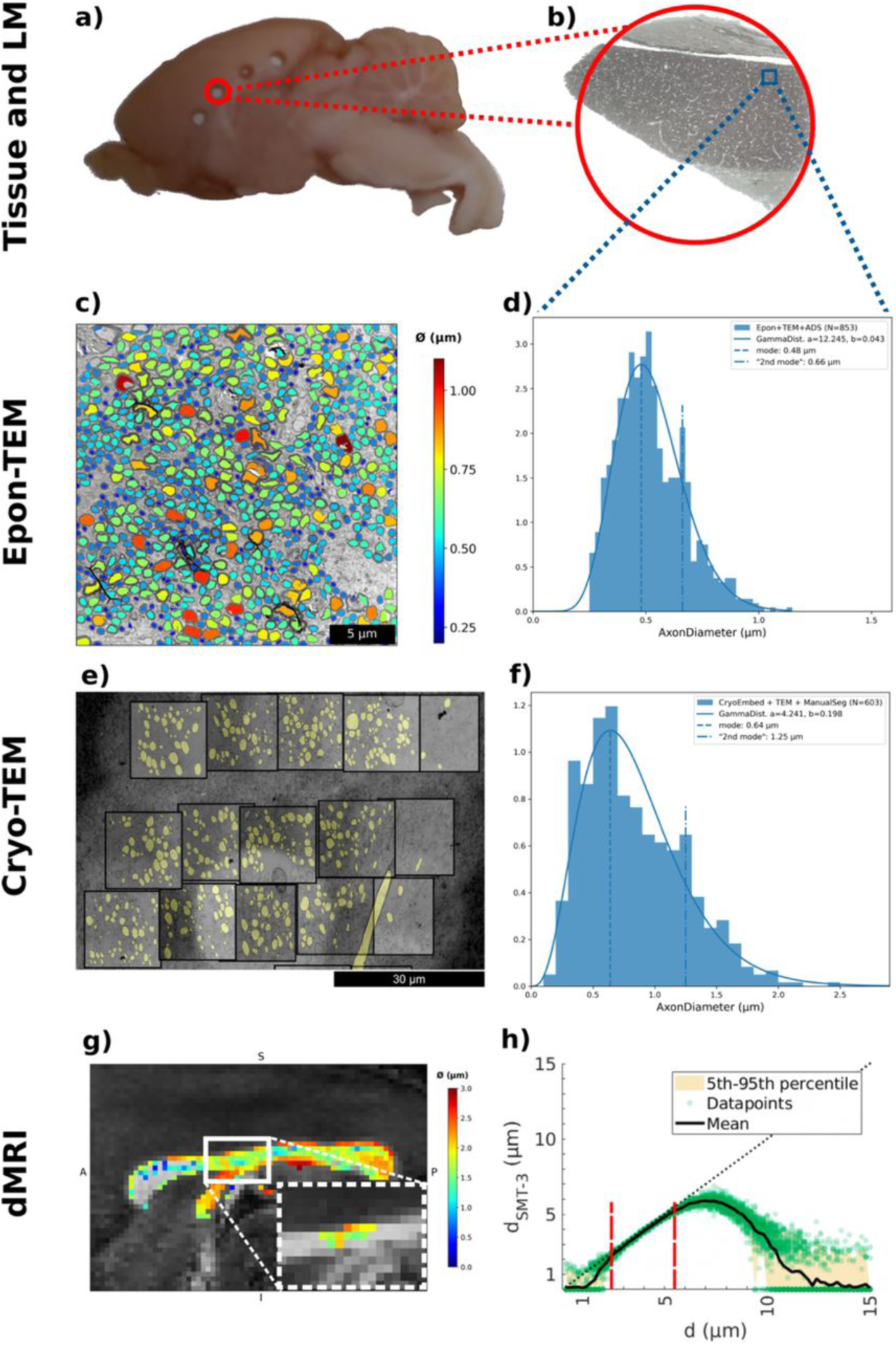
Axon diameter measurements in corpus callosum using different imaging techniques. **a)** Mid-sagittal slab (∼ 3 mm thick) with Ø = 1 mm punctures. **b)** Cross section of the EPON-embedded puncture, imaged with light microscopy at 10x magnification. **c)** 4200x magnification sample window from rat14.1 (same as in panel g), roughly corresponding to the blue square insert in panel b. Segmentation of axons is colored according to their diameter (color bar to the right). **d)** Histogram of the distribution of axon diameters in CC automatically segmented and estimated with AxonDeepSeg of this particular TEM image in panel c, from the CC in the Epon-embedded and sectioned sample. **e)** Overview TEM image from Cryo-fixated and -embedded CC-tissue (Cryo-TEM) from a different animal (rat4.2). FOV: [75 x 75] µm, magnified to FOV: [19x19] µm (black box inserts) for quantification), **f)** Axon diameter distribution from TEM of the cryo-fixed and EPON-embedded CC tissue. **g)** dMRI axon diameter distribution from rat14.1 (same as in panel b) of whole CC.A = anterior, P = posterior, S = superior, I = inferior. The white box insert indicates the ROI voxels determined by tractography from the same rat. **h)** The sensitivity profile to the diameter of the dMRI acquisition for Rician noise with a signal-to-noise ratio of 50, showing the estimated cylinder diameter vs. the ground truth diameter, d.

The axon diameter distribution associated with Epon-TEM tended to have a slightly visible shoulder (or “second mode”) of 0.66 µm in the representative image case (Fig. 3d, dash-dotted line). The mean shoulder for this animal (across 79 TEM images) was 0.49 µm, and was apparent in all animals with a group mean (± SD) of 0.51 ± 0.08 µm (N=5; Supplementary Fig. 2). The apparent second mode indicates a possible bimodal distribution of the myelinated axons within the CC, corresponding to smaller and larger diameter sub-populations.

To explore how tissue shrinkage of classical TEM sample preparation and embedding (Epon-TEM) affected the shape of the axon diameter distribution compared to the *in vivo*-like state, we performed cryo-fixation and -embedding of tissue from another similar animal (rat4.2). In the same CC region as for Epon-TEM, we manually segmented axons (N=603) from the 15 collected Cryo-TEM images, as shown in Fig. 3e. The fitted gamma distribution (a = 4.341, b = 0.194) has a mode of 0.64 µm (mean = 0.84 µm; var = 0.16; skew = 0.96), which is one 33% higher than the mode for the Epon-TEM. Compared with Epon-TEM, the tail of the distribution was wider, having a skewness of 0.96 compared with 0.57 for the representative Epon-TEM image. The Cryo-TEM histogram showed a “shoulder”, as in Epon-TEM, in the axon diameter distribution at 1.25 µm (Fig. 3d, dash-dotted line). This shoulder was more clearly separated from the primary mode in comparison to Epon-TEM (Fig. 3d vs. 3f) and is 48% higher than in Epon-TEM. The larger axons belonging to the second mode shrink more than those belonging to the first and dominating mode.

The g-ratios were estimated from the TEM segmented axons (Supplementary Fig. 1), which produced a group mean (± SD) of the g-ratio distribution medians of 0.68 ± 0.03 (N=5). Axon diameter distributions as well as g-ratio distributions overlapped between animals (Supplementary Fig. 1 and 2).

Using dMRI we estimated the diameter along the bilateral M1 projection within individual trajectory ROIs delineated by tractography between the stimulating and recording regions (Fig. 2c). Figure 3g shows axon diameter estimates in a midsagittal slice of the CC, and an outline of the tractography ROI where M1 projects through. Across the CC, as seen in Fig. 3g, as well as along the contralateral M1 tract, some voxels show axon diameters of zero. Excluding these voxels along the M1 projection, the distribution mode of this animal (rat14.1) was 1.34 ± 0.42 µm (mode ± SD). For the voxel distributions of all animals, see Supplementary Fig.3. Fig. 3h shows the simulated expected sensitivity profile of the experimental dMRI setup to axon diameter when accounting for the acquisition parameters and SNR[28]. There were lower and upper diameter bounds of the linear sensitivity range around 2 and 5.5 µm, respectively. Below the lower bound, there was a broader transition area between 1 - 2 µm with large uncertainty in diameter estimations (Fig. 3h). Hence, MRI voxels in which the diameter is estimated to be zero could potentially correspond to real diameters of 2 µm or lower.

### 2.4. Correlative Multi-modal structure-function correlation

We map the structure-function correlation between the axon diameter distributions from the different structural image modalities (dMRI, TEM, Cryo-EM) and functional measure (TCT), as well as their corresponding predicted counterparts as shown in Fig 4. The conversion between the measured and the predicted TCTs and axon diameters use Eq. 1 and Eq. 2 in Methods and is based on the group mean g-ratio (0.68 ± 0.03; N=5) and the group mean pathway length (11.47 mm ± 0.47, N=8).

**Figure 4:**
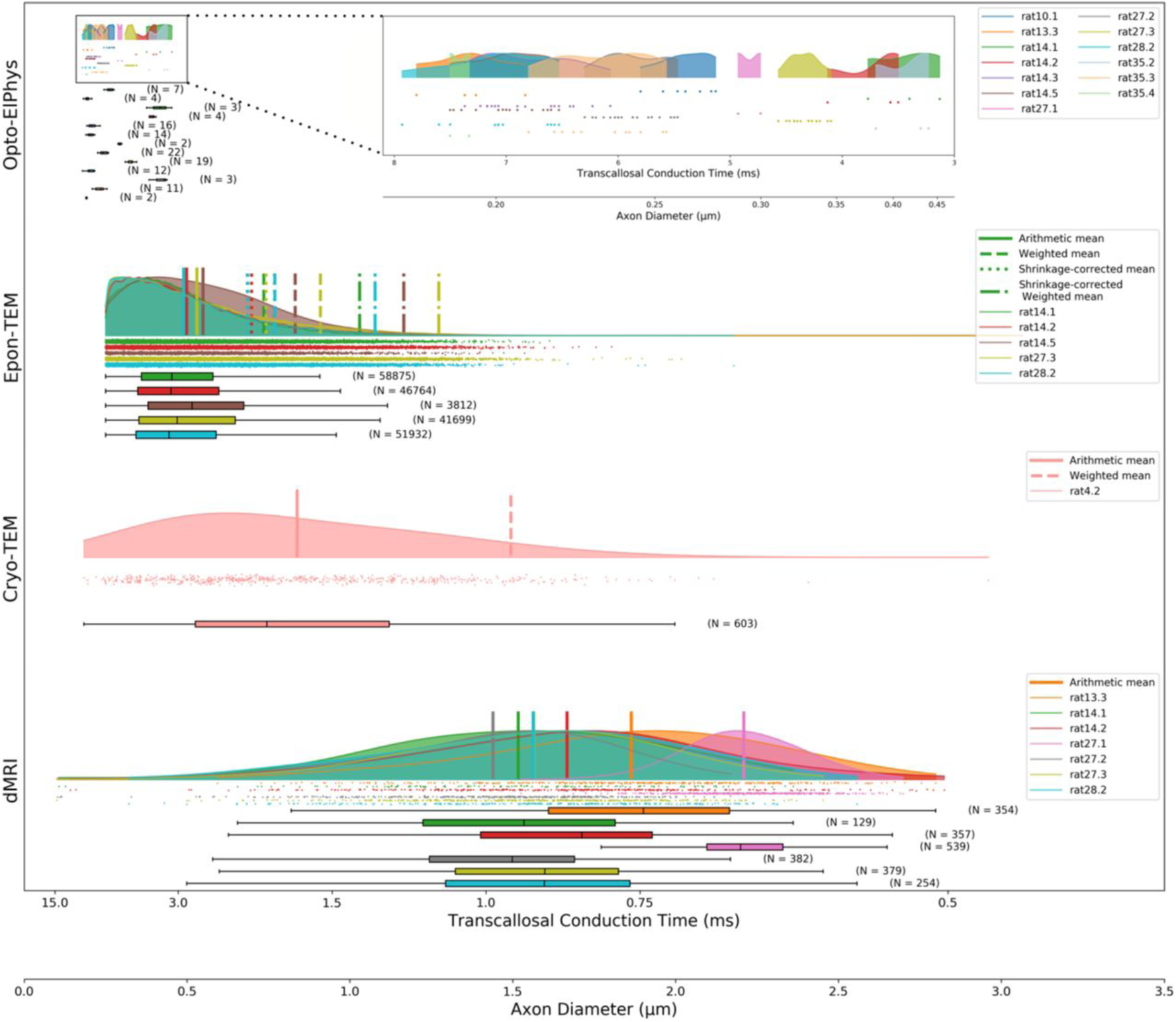
Structure-function relationships of measured and predicted values of all the modalities applied to the rat brain. Data is presented on a double x-axis, which represents latencies and diameters, respectively. Hence, individual points lying below the half-violin correspond to individually measured data points (denoted “N”) of either the transcallosal conduction time in ms (Opto-ElPhys) or axon diameters in µm (Epon-TEM, Cryo-TEM, dMRI). Each distribution corresponds to data from one animal (depicted in different colors). For the axon diameter distributions (TEM and dMRI data), the arithmetic means are shown as solid vertical lines. For the TEM data, the weighted mean diameters (for comparison with dMRI) are shown as dashed vertical lines. For the Epon-TEM data, the mean of the shrinkage-corrected distribution as well as the weighted mean diameter are shown as dotted vertical lines and dash-dotted lines, respectively. The two x-axes depict the corresponding prediction of TCT, based on axon diameter, and vice versa - converted by a fixed g-ratio (g = 0.68; N = 5 rats) and the mean estimated projection pathway from tractography of all dMRI scanned animals (L = 11.47 ± 0.47 mm; N = 8 rats).

The prediction of the mean axon diameter from the group mean (± SD) of TCTs (5.57 ± 1.49 ms; N=13) corresponded to the axons with mean (± SD) diameters of 0.28 ± 0.09 µm (N=13). Such diameters extended from below and up to the mode of the diameter distributions quantified with TEM.

For shrinkage compensation of axon diameters obtained from Epon-TEM we applied a fixed mean (i.e., of 33% and 48%, Fig. 3d vs. 3f) shrinkage factor of 40% to the axon diameter distributions (N=5). Shrinkage-corrected means of the Epon-TEM axon distributions (vertical dotted lines, Fig. 4, row 2) are shifted closer and thus show good correspondence to the arithmetic means of Cryo-TEM (Fig. 4, row 3; vertical solid lines, Supplementary Fig. 3).

The predicted TCTs correspond with the population of small axons below and up to the mode of the diameter distributions from Epon-TEM and Cryo-TEM (Fig. 4, row 1 vs. rows 2 and 3).

The arithmetic mean voxel distribution of dMRI-estimated diameters shown in Fig. 4 (row 4) have no simple relation to the predicted TCT, due to the weighting of the dMRI diameter estimate towards larger diameters related to Eq. 3 (Methods). The weighted mean of the axon diameters from shrinkage-corrected Epon-TEM (dash-dotted vertical lines, Fig. 4, row 2) and weighted mean from Cryo-TEM (dashed vertical lines, Fig. 4, row 3) generally agree, but are lower than those obtained from dMRI. Note, the smaller diameter axons, likely responsible for the measured TCT in Figure 4 (row 1), are not directly quantifiable with dMRI.

The between-animal variation of the dMRI-estimated diameters is larger than the TEM results obtained on the same animals. This suggests that axon diameters detected by dMRI in some animals mostly fall within the non-linear transition region of the sensitivity profile (Fig. 3h), which likely accounts for the large between-animal variation.

## 3. Discussion

Our main finding was that the TCT from LFP measurements in the intact rat brain corresponds to the small axonal subpopulation with diameters extending up to the mode of the diameter distribution as obtained with histology. Thus, we could not correlate the structure and function relationship of the population of larger axons that dMRI is sensitive to. We found large variations in the estimated axon diameter distributions across the different modalities. However, since we used the same animals, we can attribute the variation to the employed imaging method and tissue preparation procedures that can introduce a significant axon population-dependent shrinkage factor. Hence, our findings suggest that one must consider the sensitivity profile of the measuring methodology for the correct interpretation of the predicted Hursh-based [2] structure-function relationship.

### 3.1. Predicting axon diameters from the transcallosal conduction times

We used the peak of the first positive deflection of the transcallosal response, P1, as an indication of TCT. Our latency measurements of the P1 peak were stable on a sub-millisecond scale within animals (SD: ± 0.26 ms), despite having used different stimulation parameters [36]. However, the TCTs between animals, varied by several milliseconds (SD: ± 1.49) – not explicable by the minor variation found in pathway length by tractography [20,37,38]. Previous studies approximated the transcallosal pathway as a straight or curvilinear distance between the stimulation and recording position [17,39] which results in underestimation of the pathway and thus predicts a faster TCT. Notably, none of the previous studies collected axon diameters and TCT in the same animals.

The observed functional variation between animals might instead be explicable by differences in electrode depths. To record the evoked LFP response, we used stereotrodes with a fixed implantation position. The standard deviation of the electrode depth was about ∼50 µm, i.e., 5%, of the intended depth of 1 mm in M1, as revealed with high-resolution *ex vivo* MRI. However, Iordanova and colleagues [40] demonstrated that the local voltage differences of the transcallosal evoked response in the S1-forelimb area are greatly affected by the cortical depth of the recording electrodes (cf. their Fig. 5d) [40]. This may explain the amplitudes and latencies variation of P1 and N1 we observed in the M1 area. Saiki and colleagues also performed optogenetic stimulation of M1, acquiring multi-unit activity (MUA) measurements with a high temporal resolution in the contralateral M1 ^66^. They found transcallosal responses from multiple individual neurons, with a mean (± SD) latency of 5.2 ± 1.8 ms, which generally agrees with our LFP findings. Because of the higher density of smaller axons in the CC, one might expect that the MUA recordings in the cortex would correspond to these small-diameter axons explaining the agreement with our LFP results. However, a single-unit or MUA approach would likely allow us to record and focus on the spiking also governed by the larger axons, despite their lower density - an opportunity not possible with our LFP recordings.

Despite the between animal variations in TCT, our results agree with other studies. Hoffmeyer and colleagues [41] recorded the intracortical transcallosal responses as LFPs in the *sensory* area of rats after *electrical* stimulation in the contralateral homologous area. They reported a conduction delay as brief as 4 ms, and the latency to the first peak of 9 - 11 ms after stimulation. They used a fixed electrode position between 300 - 400 µm, i.e., considerably closer to the cortical surface than our recordings. Seggie and Berry [42] performed cortical *electrical* stimulation and LFP recordings of the interhemispheric response in rats and found 8 ms latencies of the first peak [42]. They stimulated and recorded in a cortical area (±3 mm lateral to the bregma) with electrodes placed on the cortical surface. Although a more indirect way of measuring the transcallosal response, their method may have spared confounds due to differing electrode depths between animals.

In humans, investigations of the TCT between the motor cortices using a dual-coil transcranial magnetic stimulation approach have given latencies of 5-13 ms [43–45], and 8-10 ms when stimulating the dorsal premotor cortex [46]. Ni and colleagues measured interhemispheric facilitation and inhibition in 0.1 ms intervals and found that the largest proportions of fibers governing the facilitation at 3.8 ± 0.4 ms and inhibition at 9.0 ± 0.3 ms [47]. Interestingly, their results give a hint that a distribution of differently sized axons governs the transcallosal response. This agrees with our results that the P1 peak measured with LFP corresponds to an axonal subpopulation at the mode of the diameter distribution dominated by the smaller axons.

Interestingly, the TCT measured with TMS in humans, as well as measures [21] and predictions [19,20] from monkeys, are comparable to those obtained in our rat study. The relatively constant TCT across species indicates structural and/or functional adaptations since the interhemispheric pathway length increases with brain size. Various studies indicate that the largest CC axons increase with brain size [17,39,48], at least to a certain degree, which predicts faster conduction velocity in those larger axons [17,39]. However, irrespective of brain size, the modal axon diameter is relatively constant across mammalian species [17,24,49]. Minor variations around this distribution mode cannot compensate for the increased conduction distances incurred by the 750-fold greater volume of the human brain compared to the rodent brain [23,50,51]. Not even tissue shrinkage effects of on average 40% can explain the constancy of TCTs despite such differences in brain size. For example, using the TCTs estimated with TMS in humans (5-13 ms [43–45]) and an estimated interhemispheric pathway length (111.4mm[20]-122.4 mm[17] for the motor sector), and a g-ratio of 0.7, Eq. (1) and (2) (Methods) predicts axon diameters in the range of 1.09 – 3.12 µm. This range corresponds to a population of axons exceeding the mode of human axon diameter in the CC [17] and is in alignment with the expectation that TMS primarily activates the larger excitatory neurons [52,53]. In contrast, using the LFP-based recordings in rats, we found that TCT corresponds to the axons of diameter below the mode. Thus, scaling of axonal diameter seems unlikely to account for TCT constancy across species of differing brain sizes. Instead, it supports our hypothesis that the axon diameter distribution includes different populations with varying latencies, suggesting distinct functional encoding contributions. It is therefore essential to consider the specificity and sensitivity profile of an applied stimulation and electrophysiological recording technique (e.g., optogenetics and contralateral LFP recordings vs. TMS and peripheral EMG recordings) when predicting the structural correlates. Similarly, when predicting conduction velocities from an axon diameter distribution, one cannot generalize to the mean or mode of the distribution (as is traditionally done) but must consider the possibility of subpopulations of differing axon diameters[54].

### 3.2. Measured axon diameter distributions depend on the tissue preparation

We observed variations in the measured axon diameter distributions depending on the tissue preparation method, which influences the prediction of conduction velocity. In the distribution estimated from Epon-TEM images, we observed a weak “shoulder” after the primary mode – or, in other terms, a second mode. This is indicative of two axonal sub-populations with smaller and larger axon diameter distributions [54]. When using cryo-fixation, which should maintain a structure closer to an *in vivo* state, we also found two modes, but with better separation between the primary and secondary modes. Interestingly, when comparing the two tissue preparation techniques, the larger axons were more affected by shrinkage than were the smaller axons, i.e., 48% versus 33%. The non-linearly appearing shrinkage differences might bear some relation to conduction velocities. In particular, internal resistance for the sodium ions within the axon affects the velocity of their movement [55]; the larger the axon, the lower the electrical resistance [56], which likely reflects lesser interference from macromolecules of the cytoskeleton. Altogether, this suggests a higher water content in the large diameter axons, which renders them more vulnerable to shrinkage during the dehydration step, used in the tissue preparation for Epon-TEM.

In white matter, tissue shrinkage factors from 0 to 65% have been reported during sample preparation [17,19,23,57] using a linear shrinkage correction. However, realizing the non-linear diameter shrinkage correction as our results suggest would require measurement or modeling of the shrinkage of individual axons which is not possible. Instead, we applied a single linear correction (Supplementary Fig. 2), being the mean of the two shrinkage factors (i.e., ∼40% - also used in previous studies [23]), gave a good correspondence with the Cryo-TEM data. This shrinkage approximation may be applicable for the axon distribution in the rat, where the distribution tail of the larger axons is small (Supplementary Fig. 2), but might be unsuitable for the larger primate brain, where the distribution tail is more pronounced [17,20].

Since *ex vivo* MRI is obtained in hydrated tissue, applying an appropriate shrinkage factor is essential for comparing axon diameters with those obtained by Epon-TEM to avoid an underestimation of true axon diameters due to the tissue processing [22,23]. Tissue preparation-dependent shrinkage in cell morphology has been reported in grey matter when compared to cryo-embedded tissue samples [34,58]. Similarly, synchrotron imaging of fixed hydrated cerebellum samples revealed the presence of larger cell bodies compared to dehydrated samples [58]. Altogether, our results suggest that a reliable shrinkage factor correction or the application of cryo-fixation and -embedding can ensure accurate structure-function predictions with minimal dependence on the tissue processing procedure.

In this study, we observed overlapping axon diameter distributions obtained with TEM between animals. This is in accord with other studies (see Supplementary Table 4) showing similar axon distribution modes and ranges between individual animals and across species when tissue preparation is performed consistently. In an investigation of different species, Olivares and colleagues [39] found the mode axon diameter to be 0.11 - 0.2 µm for smaller animals (e.g., rats) and large animals (e.g., cows and horses) [39]. However, other groups reported a larger modal axon diameter distribution in rats (≥ 0.8 µm) [24,30] indicating the possible impact of using varying tissue preparation procedures. Therefore, comparing predictions of conduction velocity from structural data across studies calls for confirmation of conduction velocity in the same animal, as in this study.

### 3.3. The structure-function relationship with dMRI

The axon diameters estimated with dMRI had distribution modes of 1.3-2.2 µm (Fig. 4, last row). For the given pathway length, this would predict latencies ≤1 ms, which we were unable to record from LFPs. We obtained only few dMRI estimates of diameters resembling those predicted from Opto-ElPhys, but these were considered to be underestimated (cf. Fig. 3h). The relationship by Hursh [2] and others [5,10] is determined by individual axons assuming constant diameters along their lengths, and with axon diameters contributing with an equal weighting towards the distribution mean. However, the diameter estimate from dMRI is different, representing a weighted mean over the entire distribution, where larger axons have a greater contribution to the estimate [22,30–33]. Moreover, our dMRI simulation results show a sensitivity profile [26] giving non-zero diameter estimates when axon diameter is within or above the transition area of the lower bound (Fig. 3h). Below this range, the diameter estimation often fails, dropping to zero or a constant value [26,28]. Some voxels from our MRI estimates returned diameters of zero, suggesting that the weighted diameter distribution present in the tissue may have been within or below the transition area, i.e., below 2 µm. This complicates the structure-function correlation of dMRI diameter estimates with conduction times from the LFP method which is unfit to detect large, fast-conducting axons; the two methods are sensitive and accurate within different diameter ranges.

Although some of the dMRI axon diameter estimates overlapped with the shrinkage-corrected and weighted Epon-TEM means, the weighted mean axon diameter from Cryo-TEM still agreed better with the dMRI estimates. We attribute this difference to the imperfect shrinkage correction we applied to the Epon-TEM axon diameters. Interestingly, dMRI showed a wide diameter distribution within animals, as also seen for Epon-TEM, in comparison to our predictions from Opto-ElPhys. Conversely, we also observed a larger between-animals variation in dMRI, as most axon estimates are found within the non-linear transition area. Some variation in the dMRI diameter estimates for rescans and across primate, subjects are however expected [26,59]. Nevertheless, even when comparing with shrinkage-corrected Epon-TEM diameters, we did not detect many axons of diameter exceeding the lower bound. We covered a large field of view (∼10,000 µm^2^ for N=1; 42,000-47,000 µm^2^ for N=4) with Epon-TEM at 4200x magnification, thereby including most of the CC contained in the 1 mm puncture sample (see Supplemental Figure 4). This was done to also capture the sparsely distributed large axons in the CC, to accommodate a methodological consideration mentioned in previous studies - namely a potential bias towards the densely packed smaller axons [30,54,60]. Further, we have recently shown that the mean diameter acquired from a 2D slice matches the mean 3D diameter along the trajectory of individual axons [54]. Therefore, even though we captured the axon diameters in an ultra-thin mid-sagittal 2D slice, this should also reflect axon diameters in 3D, as probed by dMRI.

In dMRI, the diameter lower bound is set by the gradient strength, acquisition parameters, SNR, and the model used to estimate diameters [26,59]. Veraart et al. used stronger gradients of 1500 mT/m versus our 600 mT/m and thereby reduced the diameter lower bound to about 1.6 µm [30]. In rats examined *ex vivo*, they reported a weighted mean diameter of around 2.5 µm in the CC region of M1 projections. This agrees with our finding of 2 µm, and supports the simulation results (Fig. 3h). that detected axons are within the transition area, explaining the larger variation when compared with shrinkage-corrected Epon-TEM of the same animal. Studies have presented *in vivo* axon diameter estimations in the human brain [22,30,59,61] with robust rescan findings [62]. Due to the hardware and SNR, those dMRI studies also had a diameter lower bound on a scale of several micrometers [22,28]. Hence, the dMRI axon diameter estimates in humans should agree with TMS-based latencies, but not with latencies from the slower-conducting smaller axons. Further, the fitting failed in some voxels of the rat brain, whereas whole-brain axon maps are obtained for the human brain [30,59]. This discrepancy supports the existence of a population of much larger axons throughout the white matter of the human brain, which is absent from the smaller rat brain.

### 3.4. Considerations

We used optogenetics to stimulate specifically the excitatory callosal projecting neurons. This is in contrast to electrical stimulation, where all neuron types in the vicinity of the electrode tip are affected [63]. Indeed, our histological investigation showed that the marker EYFP was profusely expressed in the CC projection area [36], confirming the preponderance of excitatory projection neurons. However, given that the callosal projecting cortical neurons are embedded in a matrix of excitatory pyramidal cells and inhibitory interneurons [64,65], some off-target neurons might also have been virally transfected.

Furthermore, the proportion of the segmented axons from TEM that were also optogenetically stimulated is unknown. Since only a very few axons in the CC are GABAergic [66], they would likely not have markedly affected the segmented axon diameter distribution or the LFP signal. The different neurotransmitter components (glutamate and GABA) of the transcallosal evoked response have been investigated with pharmacological challenges during electrical [41,67] and optogenetic stimulation [68,69]. Results of those studies indicated that the initial response (P1) is likely the excitatory postsynaptic potentials caused by the transcallosal glutamatergic response, whereas the subsequent peak (N1) might be inhibitory postsynaptic potentials from the local inhibitory GABAergic interneurons.

Although optogenetics is focally specific to neuron type ^67^, and should not introduce a stimulation artifact, it has a potential caveat concerning temporal delays in comparison to electrical stimulation. The temporal delay (t_kON_) of hChR2-H134R is between 0.6 ms and 2.6 ms ^68–70^ depending on the stimulation intensity. As the latencies obtained from the optogenetic experiment arose from different laser stimulation intensities, adjusting for the onset latency would require individual adjustment values. Furthermore, with our experimental setup, we could not measure the actual optogenetic stimulation delay, which would have required a patch-clamped approach with an additional electrode at the stimulation position. Applying a single fixed stimulation intensity would likely have given rise to even smaller differences in the obtained TCT. That said, we obtained a sub-millisecond variation of latencies within animals, which suggests that the range of stimulation intensity had a minimal differential effect on the latency. Taking into account the onset latency, the TCTs would likely have overlapped with a broader distribution of the axon diameters, including axons larger than the mode of the axon diameter distribution. This indicates that the prediction of axon diameters based on TCT should in the future consider the temporal delay of the onset kinetics for optogenetic activation.

## 4. Materials and Methods

Below is a summarized version of the materials and methods. Further details can be found in the Supplementary Information.

All animal procedures were conducted in accordance with the ARRIVE guidelines, the European Communities Council Directive (2010/63/EU) and were approved by the Animal Experiments Inspectorate (2016-15-0201-00868) of Denmark. The electrophysiological raw data were collected as part of our previous study [36], where detailed methods can be found. The data was reanalyzed in the present study to obtain the TCT.

### 4.1. Animals and surgery

16 young male Sprague-Dawley rats (NTac:SD-M; 4 weeks old) underwent stereotaxic surgeries. We made craniotomies above the bilateral M1 (AP: +1.0 mm; ML: ±2.5 mm, relative to bregma). In the right M1, we injected the viral inoculum (AAV5::CamKIIα−hChR2(H134R)−EYFP; UNC Vector core) at a depth of -1.00 mm relative to the dura followed by implantation of a calibrated optic fiber implant (Ø = 55 µm). In the left M1, a stainless steel stereotrode pair (Ø = 127 µm) was implanted at -1.00 relative to the dura, to record the evoked potentials as LFPs.

### 4.2. Electrophysiological recording of optogenetically evoked transcallosal potentials

The animals were exposed to laser light stimulation (𝛌 = 447 nm) while recording the evoked responses as LFPs during dexmedetomidine and low dose isoflurane anesthesia. For peak detection, all trials for each condition were averaged. The TCT was represented by the latencies of the first measurable transcallosal response (P1). Data from two animals was excluded for lacking reliable evoked potentials.

### 4.3. Perfusion fixation

The rats were perfusion fixed immediately after the final experiment. Transcardial perfusion was initiated with 0.1 M potassium phosphate-buffered saline (KPBS) delivered at ∼15 mL/min for approximately 3 minutes, followed by 7-12 minutes of infusion with 4% formaldehyde. The brain was extracted and post-fixed at 4 °C for at least 3 weeks.

### 4.4. Post mortem MRI

#### 4.4.1. Preparation prior to MRI scan

At least three weeks before MRI scanning, a subset of the brains (N=9), were placed in 0.1 M KPBS [23]. The brains were scanned in a double lined plastic bag containing a minimal volume of room temperature KPBS, on a sample holder, custom-made with LEGO^TM^ [51]. The samples were scanned in a cryo-coil setup on a 7T preclinical Bruker scanner (Bruker BioSpec 70/20 USR) using Paravision 6.0.1.

#### 4.4.2. Diffusion MRI

For axon diameter estimation, we used a three-shell pulsed gradient spin echo (PGSE) protocol with single-line readout. Three unique b-values with b = [24.450, 21.247, 17.560] s/mm^2^, gradient strengths G = [590, 550, 500] mT/m, gradient separation (Δ) = 18 ms, and gradient duration (∂) = 8 ms were applied along 30 uniformly distributed diffusion gradient directions [70,71]. In addition, nine b=0 images were acquired. All shells used the same echo time (TE) of 28.5 ms and repetition time (TR) of 3500 ms and were acquired with an isotropic voxel size of 125 µm. Ten sagittal slices with no slice gap covered a limited field of view around the midsagittal region of the CC. Total acquisition time was 35 hours.

For tractography, whole-brain dMRI data sets (isotropic voxel size of 125 µm; 80 axial slices; no slice gap; matrix: 128 x 128 x 80; FOV: 16mm x 16mm x 10 mm) were collected using a single shell with b-value of 4000 s/mm^2^ (gradient strength (G) = 224 mT/m; Δ = 23 ms; ∂ = 7.5 ms) in 61 isotropic distributed non-collinear directions [72]. The scan was acquired with TE = 38.6 ms and TR = 3592 ms. Total acquisition time was 9 hours.

To assess the lower bound of measurable axon diameter, we calculated the sensitivity profile to axon diameter of this dMRI acquisition [28].

The dMRI data sets were denoised using the variance stabilizing transform and optimal shrinkage singular value manipulation method [73], and processed to remove Gibbs ringing artifacts [74] using the MRTrix3 software toolbox (RRID: SCR_006971).

#### 4.4.3. High resolution structural image

A high-resolution structural 3D T2 weighted MR image was obtained for five rats for determination of implant depths. A “true” fast imaging with steady-state free precession (FISP) T2 weighted sequence acquired high-resolution whole-brain visualization with the following parameters: TR: 2.5 s; TE: 5.1 ms; matrix size: 256 × 256 × 128; field of view: 23.04 × 23.04 × 11.52 mm^3^; Image resolution: 90 × 90 × 90 µm^3^; Flip angle: 30 degrees, Averages: 40; and total acquisition time: 2 hours.

#### 4.4.4. Tractography

We performed probabilistic streamline tractography on the whole brain dMRI data set. The tract was a summed length of two tractograms, performed for each hemisphere. Seed ROIs were placed below the optical fiber or electrode depth, and set to terminate in the midsagittal CC. Up to one million streamlines were seeded using the IFOD1 function in the MRtrix3 toolbox ^50^, with selection of a least 1000 streamlines. One dataset was excluded due to unsuccessful tracking.

#### 4.4.5. Axon diameter estimation from dMRI

The preprocessed multi-shell dMRI data sets were normalized by the voxel-wise average of the b=0 images. The powder average for each b-value was calculated via the arithmetic average of the signals across the 30 diffusion encoding directions. The extra-axonal space could be approximated to be fully attenuated due to the high b-values [30,75] and the signal was assumed to only contain contributions from the intra-axonal space. An axonal signal model describing the spherical average of cylinders [54,59,76,77] was fitted to the powder average signals using the non-linear least squares [54].

Three unknown parameters of the following variables were fitted: the intrinsic diffusivity i.e., intra-axonal axial diffusivity (D_o_), the intra-axonal perpendicular diffusivity (𝐷_⊥_), and the signal fraction of the intra-axonal space (𝑣_*a*_). The diameter could then be calculated from 𝐷_⊥_ [78]. The intrinsic diffusivity was measured in individual brains from the principal direction within the CC region using the diffusion-tensor model fitted to the b-value of 4000 s/mm^2^, and was kept constant for the fitting of axon diameter [22].

Finally, we estimated the axon diameter from dMRI in the bilateral M1 projection region of CC. A 3D ROI was placed in the projection region of CC automatically mapped by the streamlines from tractography. Axon diameters below 0.1 µm were discarded.

### 4.5. Histology

A subset of the brains was manually sectioned into two hemisphere slabs (3-4 mm thick).

#### 4.5.1. Fluorescent LM - locating the transcallosal fibers of M1

The right hemisphere was sectioned sagittally into 40 µm-thick slices. A “super image” was obtained using a series of 10x magnification images that were stitched. Inspection of the sagittal slice revealed enhanced yellow fluorescent protein (EYFP) expression in the dorsal-anterior midbody of CC and informed the subsequent puncture procedure.

#### 4.5.2. Histological preparation TEM

##### 4.5.2.1. Epon embedding of fixed tissue

The midsagittal slab from the left hemisphere was punctured using a Ø = 1mm biopsy puncture tool covering the region corresponding to projection locus of CC. The samples were embedded in 2% Agar and transferred to a 2.5% solution of glutaraldehyde in cacodylate buffer (0.1 M) and stored for two weeks. Thereafter, Epon embedding was conducted following a standard procedure [18].

##### 4.5.2.2. Cryofixation of fresh tissue

One rat (NTac:SD-M) was perfusion fixed with minor adjustments. The perfusion with 4% formaldehyde was carried out for only 3 minutes at 15 mL/min. We obtained 200 µm-thick slices of the midsagittal slab, with a similar region of the CC as above was sampled using a biopsy puncture. Following two hours of storage in KPBS, the sample underwent high-pressure freezing (HPM100; Leica Microsystems, Wetzlar, Germany), immediately followed by freeze substitution (AFS2; Leica Microsystems, Wetzlar, Germany) using 1% OsO4 in acetone, and consecutive Epon infiltration and embedding of the sample. Ultrathin sections (50-70 nm) were made with an ultra-microtome. Contrast enhancement was performed using a double staining protocol with uranyl acetate and lead citrate solutions (EM AC20; Leica Microsystems, Wetzlar, Germany).

##### 4.5.2.3. Sectioning for TEM

After coarse trimming of the blocks we obtained ultra-thin sections (∼ 40-70 nm) using an ultramicrotome (Ultracut UCT or EM UC7, Leica Microsystems, Wetzlar, Germany). The sections were automatically stained on the grids with uranyl acetate and lead citrate (EM AC20; Leica Microsystems, Wetzlar, Germany).

##### 4.5.2.4. TEM of Epon-embedded samples

TEM images were obtained on a CM100 microscope (Philips; FEI, Eindhoven, NL) at 4200x magnification, captured in a sequential manner, to cover the entire CC cross section in the Ø=1 mm punctured sample. This resulted in between 16 images from the dorsal part of CC (N=1) to 69-77 images (N=4) at an image FOV of ∼25 x 25 µm. The segmentation of the acquired images was obtained using AxonDeepSeg [35]. The model provided individual inner (d) axon diameters and corresponding g-ratios of all segmented axons. The acquired axon diameter distributions were filtered between 0.25 and 5.0 µm, based on visual inspection.

##### 4.5.2.5. TEM of Cryo-fixed tissue samples

TEM images of the cryo-fixed were obtained on the same microscope, with an image FOV of 17.5 x 17.5 µm. Fifteen TEM images were obtained from different positions, within the same section - and 607 axons were manually segmented in ITK Snap (RRID:SCR_002010). A connected components analysis of the segmented axons was performed in MATLAB 2020a. Owing to the eccentricity of many of the axonal cross sections, the minor axis of an ellipse was used to represent axon diameter. The obtained axon diameter distributions were filtered between 0.1 µm and 5.0 µm, resulting in 603 axons.

### 4.6. Calculating the structure-function relationship between conduction velocity and axon diameter

From the obtained length of the projection pathway (L) and the measured TCT, we calculated the estimated conduction velocity (CV) as follows:

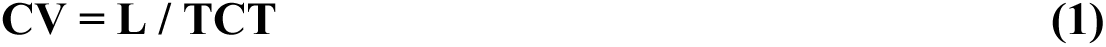

Similarly, we calculated the conduction velocity along the pathways from the measured g-ratio and axon diameter, given Hursh’s structure-function relationship [2,5,10]:

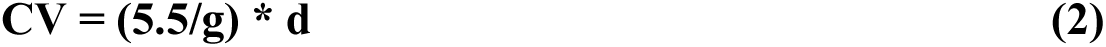

where g, i.e., the g-ratio, was set to group mean of TEM data results, rather than the literature value of ∼0.7 [10,20,54]. Hence, combining Eq. (1) and (2) allows us to predict the TCT from structural measures, or the converse prediction of axon diameter from the functional measure.

### 4.7. Statistical Analysis

#### 4.7.1. Axon diameter distributions from histology

We fitted gamma distributions with parameters *a* (shape), and *b* (scale), and used a location parameter of 0. Statistical features were expressed as the mean, SD, and the mode. The voxel-wise dMRI diameter estimates are weighted towards larger diameters [22,30–33]. We accounted for this weighting by calculating the weighted diameter (*d_w_*) from the histological axon diameter distributions. In the wide pulse limit, 𝛿 ≫ 𝑅^2^⁄𝐷_0_[30,32,33], where *R* is the axon radius from histology, *D_0_* is the intrinsic diffusivity and 𝛿 is the pulse-width of the diffusion encoding. In practice, this wide-pulse limit applies to most axons in the rat brain [28] Using this assumption we can calculate *d_w_* as [30,32]:

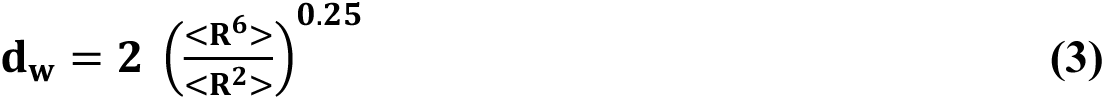

## Acknowledgments

Thanks to local Laboratory Technician, Sascha Gude, for her immense help with animal caretaking and assisting in experiments, throughout the project period. Thanks to laboratory technician Susanne Sørensen from Bispebjerg Hospital for her help with sample preparation for both light and electron microscopy. Thanks to H. Martin Kjer for aiding in initial axon segmentation methods for LM data - and ideas for visualization. Last, but certainly not least, thanks to the late Bente Pakkenberg for her invaluable inputs and guidance during laboratory experiments and manuscript revision – initially as a co-author. The present study builds upon her pioneering work on corpus callosum and she will be sincerely missed as a person - and researcher.

## Author Contributions

Conceptualization: CSS, HRS and TBD; Methodology: CSS, MA, MP and TBD; Investigation: CSS, MA, TBD; Visualization: CSS, MA; Writing–Original Draft: CSS, TBD; Writing–Review & Editing: CSS, MA, MP, HRS, TBD; Funding Acquisition: HRS and TBD; Resources: HRS, TBD; Supervision: HRS, TBD.

