## Supplementary Information for "Mapping axon diameters and conduction velocity in the rat brain – different methods tell different stories of the structure-function relationship"

**This PDF file includes:**

Supplementary Materials and Methods
Figures S1 to S4
Tables S1 to S4
Supplementary Information References

**Supplementary Materials and Methods**

1. Experimental Design

The objective of this study was to relate measurements of axonal structure (diameter and myelin) with measurements of their function (conduction velocity), uniquely from the same animals. The transcallosal conduction time between the bilateral motor cortices was measured as LFPs with electrophysiology after contralateral optogenetic stimulation. The conduction pathway was measured with tractography from ex vivo dMRI, to calculate the conduction. velocity. The axon diameter was initially estimated with non-invasive *ex vivo* dMRI and subsequently with TEM. To investigate shrinkage caused by the classical tissue processing for TEM, we also acquired Cryo-TEM of the same region in one animal.

The electrophysiological raw data were collected as part of our previous study, investigating the reliability of the evoked LFPs with varying optogenetic stimulation parameters [1]. The functional metrics, latency of the first transcallosal response, is shown only in the present study. All animal procedures were conducted in accordance with the ARRIVE guidelines, the European Communities Council Directive (2010/63/EU) and were approved by the Animal Experiments Inspectorate (2016-15-0201-00868) of Denmark.

**2. Animals and surgery**

16 young male Sprague-Dawley rats (NTac:SD-M; 4 weeks old) underwent stereotaxic surgeries (Stoelting Lab Standard, Digital Dual Manipulator; Wood Dale, IL, USA) [1]. In brief, we made craniotomies above the bilateral M1 (AP: +1.0 mm; ML: ±2.5 mm, relative to bregma), where the dura was punctured. In the right M1, we injected the viral inoculum (AAV5::CamKIIα−hChR2(H134R)−EYFP; UNC Vector core) at a depth of -1.00 mm relative to the dura. A calibrated optic fiber implant (Ø = 55 µm) was later implanted at the same position. In the left M1, a stainless steel stereotrode pair (Ø = 127 µm) was implanted at the same depth to record the evoked potentials as LFP. The signal was amplified with gain factors of 100x - 1000x. In the former case, we multiplied the recorded signal by a factor of 10. Hardware filters were set to bandpass between 0.1 Hz and 50 kHz. The recorded LFP data was further filtered with a notch filter at 50 Hz and a 2nd order Butterworth band-pass filter between 3 Hz and 300 Hz. For further details, see the previous study [1].

**3. Electrophysiological recording of optogenetically evoked transcallosal potentials**

Four to seventeen weeks after surgery, the animals were exposed to a paradigm of blue laser light brain stimulation ($\lambda$ = 447 nm) while recording the evoked responses as LFPs. Prior to the experiment, the animals had been anesthetized with dexmedetomidine and low dose isoflurane, as described in our previous study [1]. The available data from [1] included systematically varied stimulation durations and intensities (0.1-10.0 ms and 0.25-10 mW), resulting in a stimulation input parameter map of 42 different conditions. Each condition produced 150 individual trials, totaling 6300 stimuli per animal, with a few (< 8 trials per condition) being discarded for technical reasons. Trials were baseline-corrected using the mean value of the data points in the 10 ms period preceding the laser stimulation onset. The signal recorded from the two channels of the stereotrode was averaged for all trials for each individual condition. Data from one animal (rat33.1) were discarded due to lacking response after optogenetic stimulation, leaving 15 animals for peak detection and quantification.

Based on the data so obtained, we estimated the transcallosal conduction time (TCT), represented by the latencies of the first measurable transcallosal response. After visual inspection of the typical stimulation responses, we defined criteria for automatic peak detection for the first positive (P1) and negative (N1) deflections. Specifically, the time span for the two peaks was restricted to the ranges t = +1 ms to t = +9 ms for P1, and t = +5 ms to t = +20 ms for N1. The peaks were automatically detected using the following procedure: If the signal amplitude extended beyond the threshold of one standard deviation, based on the 100 ms epoc preceding the stimulus onset (from t = -105 ms to t = -5 ms), the maximum absolute amplitude of the deflection within the timespan was considered a peak. We manually inspected the results of automatic peak detection and and discarded erroneous cases. The peak onset of N1 was interpolated, as in the previous literature [2,3]. Specifically, we took the onset as the intersection of a first-order regression line on the peak slope, between 45% and 55% of the N1 peak minimum, and the baseline. For each condition, the signal amplitude of each individual trial was captured exactly at the obtained latency of the N1 peak to calculate the coefficient of variation (SD/Mean). This coefficient of variation served to determine the reliability of the peak within each condition. Any conditions providing a coefficient of variation of the N1 peak below 1.00 were considered to be reliable. One animal (rat9.4; in addition to rat33.1, mentioned above) produced no reliable conditions based on this criterion, despite showing visible peaks in some conditions. Therefore, we obtained latencies from 14 animals.

The P1 peaks of the measured LFP signal were often insufficiently prominent to detect the P1 peak onset. Therefore, as an approximation of the TCT we instead used the median of the P1 peak latencies from all reliable conditions of each animal.

The data analysis was accomplished with custom-made python-scripts (https://git.drcmr.dk/cskoven/elphys), utilizing the open source analysis toolkit for Open-Ephys-data (https://github.com/open-ephys/analysis-tools).

### 4. Perfusion fixation

To extract the brains for further *post mortem* MRI and histological analysis, the rats were perfusion fixed immediately after the final optogenetic brain stimulation session. Here, anesthesia was induced with 5% isoflurane, followed by subcutaneous injection with mixture of Hypnorm® (Fentanylcitrat 0.315 mg/mL and Fluanisone 10 mg/mL; Department of Experimental Medicine, Panum, Copenhagen, DK), sterile water (Fresenius Kabi, Uppsala, Sweden) and Dormicum® (Midazolam 5 mg/mL; B. Braun, Germany) in a 1:2:1 ratio (3 mL/kg). Transcardial perfusion was initiated with 0.1 M potassium phosphate-buffered saline (KPBS) delivered at 15-20 mL/min for approximately 3 minutes, followed by 7-12 minutes of infusion with 4% formaldehyde (CellPath; Newtown, Powys, UK). The brain was extracted and post-fixed pin 4% formaldehyde at 4 $^{o}$C for at least 3 weeks. At least three weeks before MRI scanning the brains were placed in 0.1 M KPBS to remove free formaldehyde, thereby restoring T2 relaxation and thus increasing SNR in the subsequent MRI scans [4].

### 5. *Post mortem* MRI

#### 5.1. Preparation prior to MRI scan

Before the histological process, a subgroup (N=8) of the *ex vivo* brains were MR scanned. The extracted intact brain was wrapped in a thin cloth sheet (Nonwoven swabs, Selefa; OneMed, Danderyd, Sweden), placed in a double plastic bag containing a minimal volume of KPBS, and allowed to reach room temperature prior to scanning. The brain was positioned on a mechanically stable sample holder custom-made with LEGO™ blocks to avoid introducing short-term instabilities into the acquired dw-MRI data set [5]. We used a cryo-coil setup, allowing for a 2.5-5x increase in SNR compared to regular room-temperature coils. The temperature at the surface of the non-cryo part of the cryo-coil was set to 24 degrees to ensure a constant temperature environment of the tissue during the scanning session. All MRI datasets were acquired on a 7T preclinical Bruker scanner (Bruker BioSpec 70/20 USR) using Paravision 6.0.1.

#### *5.2. Diffusion MRI*

For axon diameter estimation we used a three-shell pulsed gradient spin echo (PGSE) protocol with single-line readout. Three unique b-values with b = [24.450, 21.247, 17.560] s/mm$^2$, gradient strengths G = [590, 550, 500] mT/m, gradient separation ($\Delta$) = 18 ms, and gradient duration ($\partial$) = 8 ms were applied along 30 uniformly distributed diffusion gradient directions that were generated using the electrostatic repulsion method [6,7]. In addition, nine b=0 images were acquired. All shells used the same echo time (TE) of 28.5 ms and repetition time (TR) of 3500 ms and were acquired with an isotropic voxel size of 125 µm. Ten sagittal slices with no in-between slice gap covered a limited field of view around the midsagittal region of the CC. Total acquisition time was 35 hours.

To assess the lower bound of measurable axon diameter, we calculated the sensitivity profile to axon diameter of this dMRI acquisition as previously described [8]. The signals from cylinders were analytically generated for diameters in the range 0.2 - 15.0 µm at 0.2 µm steps. Rician noise of a given signal-to-noise ratio (SNR) was added to the signals and the powder average cylinder model was fitted to obtain a diameter estimate. The Rician distributed noise was emulated by calculating the magnitude of complex Gaussian distributed noise in which the standard deviations of the real and imaginary components were both 1/SNR. The diameter estimation was performed 50 times for each diameter, with each repeat represented by a green data point in Fig. 3h. The sensitivity profile revealed a diameter lower bound of around 2 μm and an upper bound of around 6 μm for accurate estimation of the diameter.

For tractography, whole-brain dMRI data sets (isotropic voxel size of 125 μm; 80 axial slices; no slice gap; matrix: 128 x 128 x 80; FOV: 16mm x 16mm x 10 mm) were collected using a single shell with b-value of 4000 s/mm$^2$ (gradient strength (G) = 224 mT/m; $\Delta$ = 23 ms; $\partial$ = 7.5 ms) in 61 isotropic distributed non-collinear directions [9]. The scan was acquired with TE = 38.6 ms and TR = 3592 ms. Total acquisition time was 9 hours.

After acquisition, the dMRI data sets both for axon diameter estimation and tractography were denoised using the variance stabilizing transform and optimal shrinkage singular value manipulation method [10], and processed to remove Gibbs ringing artifacts [11] using the MRTrix3 software toolbox (RRID: SCR_006971).

#### *5.3. High resolution structural image*

A high-resolution structural 3D T2 weighted MR image was obtained for a subgroup of animals (N=5) to compare with histological slices, for determination of implant depths and for visualization. A "true" fast imaging with steady-state free precession (FISP) T2 weighted sequence acquired high-resolution whole-brain visualization with the following parameters: TR: 2.5 s; TE: 5.1 ms; matrix size: 256 × 256 × 128; field of view: 23.04 × 23.04 × 11.52 mm$^3$; Image resolution: 90 × 90 × 90 μm$^3$; Flip angle: 30 degrees, Averages: 40; and total acquisition time: 2 hours.

#### *5.4. Tractography*

To obtain an estimate of the length of the interhemispheric callosal M1 pathway, we performed probabilistic streamline tractography on the whole brain dMRI data set. To identify the position of injection sites, we used the high-resolution T2w-MRI scan. In combination with histological slices of the fluorescently labeled excitatory neurons, we confirmed the positions of the fiber and electrode implantations, as well as the axonal projection region through the midsagittal slice of the CC. The most robust way to segment the tract was proved to be summation of the length of two tractograms performed individually for each hemisphere. One tractogram emanated from the optical fiber depth and the other from the electrode depth, with both set to terminate in the midsagittal CC. All ROIs were drawn on fractional anisotropy (FA) maps generated in MRtrix3. The seed ROIs for each hemisphere were drawn axially just beneath the shaft lesion from fiber or electrode implantation. The target ROI was drawn covering the whole midsagittal plane of CC. Further, exclusion ROIs were drawn axially, just superior to the seed ROIs, thereby avoiding streamlines that erroneous projected into the shafts of either electrode or fiber implantations in the cortex. Finally, a set of exclusion ROIs were placed sagittally (lateral to the implant regions) as well as coronally (anterior to the implant regions), and posterior to the CC projection area of. This was done to further exclude spurious streamline trajectories that obviously deviated from the main tract. Up to one million streamlines were seeded in the generation of each tractogram, with selection of a least one thousand streamlines. The mean (± SD) length of all streamlines was calculated. The tractography was based on constrained spherical deconvolution for fiber reconstruction, with performance of probabilistic tracking using the IFOD1 function with standard settings in the MRtrix3 toolbox [50]. In some cases, the tractography was unable to cover the full pathway into the gray matter at the fiber and/or electrode position. These path discrepancies, as well as the distance between the connecting tracts in the CC, were measured manually from the FA image. Tractography was performed on nine brains, where one was excluded due to unsuccessful tracking.

#### *5.5. Axon diameter estimation from dMRI*

The preprocessed multi-shell dMRI data sets for axon diameter estimation were normalized by the voxel-wise average of the b=0 images. The powder average for each b-value was calculated via the arithmetic average of the signals across the 30 diffusion encoding directions. The high b-values were such that the signal from the extra-axonal space could be approximated to be fully attenuated [12,13]. For the model fitting, as thus assumed that the signal only contained contributions from the intra-axonal space. An axonal signal model describing the spherical average of cylinders [14–17] was fitted to the powder average signals using the non-linear least squares approach in Andersson et al. [14]. That model assumed that the Gaussian phase approximation was valid.

The model fitted three unknown parameters of the following variables: the intrinsic diffusivity i.e., intra-axonal axial diffusivity ($D_o$), the intra-axonal perpendicular diffusivity ($D_\perp$), and the signal fraction of the intra-axonal space ($v_a$). The diameter could then be calculated from $D_\perp$using the Van Gelderen et al. (1994) formulation for the PGSE signal perpendicular to a cylinder [18]. The intrinsic diffusivity was measured in individual brains from the principal direction within the CC region using the diffusion-tensor model fitted to the b-value of 4000 s/mm$^2$, and was kept constant for the fitting of axon diameter [19].

Finally, we estimated the axon diameter from dMRI in the bilateral M1 projection region of CC. A 3D ROI was placed in the projection region of CC, lying between the stimulation and recording area in the bilateral M1 areas, and automatically mapped by the streamlines from tractography. Axon diameters below 0.1 µm were discarded. The mean axon diameters from all voxels in the ROI were obtained and the mean (± SD) axon diameter was calculated for all animals.

6. Histology

After the *ex vivo* MR scanning, the brains were further processed for histological investigations. This entailed fluorescent LM for validation of the projection pathway in mid-sagittal CC slices. Further, the tissue preparation allowed regular LM for overview images and also TEM for axon diameter estimation.

#### *6.1. Fluorescent LM - locating the transcallosal fibers of M1*

The brains were manually sectioned into two hemisphere slabs (3-4 mm thick). The right hemisphere was sectioned sagittally into 40 µm-thick slices using a vibratome (VT1000s; Leica Biosystems, Nussloch, DE) and placed on SuperFrost® slides (Menzel Gläser; ThermoScientific, Braunschweig, DE), mounted with Fluoroshield (w/ DAPI, #F6057, SigmaAldrich), and covered with Coverslips (size #0, Menzel Gläser).

A “super image” was obtained using a series of 10x magnification images, which were automatically stitched together by “Stereology” (Visiopharm; Hørsholm, Denmark) microscopy software (see Fig. 2b). Inspection of the sagittal slice revealed enhanced yellow fluorescent protein (EYFP) expression in the dorsal-anterior midbody of CC and informed the subsequent puncture procedure.

#### *6.2. Histological preparation for LM and TEM*

Having visualized where the axons from the excitatory neurons in M1 projected through CC, we prepared the same brains for LM and TEM for quantification of axon diameters and myelin thickness.

##### 6.2.1. Epon embedding of fixed tissue

To determine the axon diameter distribution histologically within the same brains, the midsagittal slab from the left hemisphere was punctured using a biopsy puncture tool Ø = 1 mm (EMSdiasum, AxLab, Denmark) covering the region corresponding to the locus where virally transfected neurons are projecting through CC. To maintain correct orientation in the axonal projection direction, the samples were embedded in 2% Agar. The Agar-embedded puncture samples were transferred to a 2.5% solution of glutaraldehyde in cacodylate buffer (0.1 M) and stored for two weeks. Thereafter, Epon embedding was conducted following a standard procedure [20]. The samples were washed for 4 x 15 min in 0.1M cacodylate buffer, stained for two hours in a 2% $OsO_4$ in 0.1 M cacodylate buffer, and then washed for 2 x 30 min in 0.1 M cacodylate buffer. The samples were then placed in fresh 0.1 M cacodylate buffer for overnight storage at 4 °C. The following day, the samples were dehydrated in a series of increasing alcohol solutions [30, 50, 70, 80, 90]% for 30 minutes each, and finally in 100% ethanol for 2 x 40 minutes. To remove residual ethanol, the dehydration was followed by submersion in propylene oxide for 3 x 10 min. Two solutions (”A” and ”B”) of Epon-812 (45345, Merck) were made with different hardeners, DDSA (45346, Merck) and MNA (45347, Merck), respectively. Solution A had a w/w ratio of 47:64 and solution B a 60:50 ratio. One hour before infiltration, the two solutions were mixed in a 1:1 ratio and 7 drops of accelerator DMP-30 (45348, Merck) per 10 mL of Epon-mixture was added. The infiltration was performed with increasing Epon concentrations in Epon:Propylene Oxide mixtures [1:1 for 1 hour, 2:1 for 1 hour] and finally in pure Epon mixture overnight at 4 °C. The following day, the samples were transferred to a new pure Epon mixture and polymerized in a heating chamber (N6c, Genlab; Widnes, England) for 48 hours at 60 $^o$C.

#### 6.2.2. Cryofixation of fresh tissue

As the classical Epon-embedding is known to cause tissue shrinkage [4,14,21,22], we also performed cryo-embedding of CC samples of fresh tissue of a similar animal neither being MRI scanned nor being exposed to the optogenetic stimulation experiment. One (rat4.2) young adult (58 days old) Sprague-Dawley male rat (NTac:SD-M) was perfusion fixed with minor adjustments from the method described above. To minimize the degree of fixation, the transcardial perfusion with 4% formaldehyde was carried out for only 3 minutes at 15 mL/min. Thereafter, the brain was carefully extracted and a sagittal slab covering the midbrain was cut using a matrix (brain matrix, EMS diasum). We then obtained 200 µm-thick slices of the slab with a Vibratome (VT1000s; Leica Biosystems, Nussloch, DE). A biopsy puncture from the central midbody of the CC was extracted with a biopsy puncture tool (EMSdiasum, AxLab, Denmark) Following 2 hours of storage in KPBS, the puncture sample was placed in an aluminum specimen carrier (Ø = 3 mm; Leica Microsystems, Wetzlar, Germany) and underwent freezing in a high-pressure freezer (HPM100; Leica Microsystems, Wetzlar, Germany). This was immediately followed by freeze substitution (AFS2; Leica Microsystems, Wetzlar, Germany) using 1% $OsO_4$ in acetone, which allowed for consecutive Epon infiltration and embedding of the sample. Ultrathin sections (50-70 nm) were made with the ultra-microtome (EM UC7; Leica Microsystems, Wetzlar, Germany) and collected on copper grids. Contrast enhancement was performed using a double staining protocol with uranyl acetate and lead citrate solutions (EM AC20; Leica Microsystems, Wetzlar, Germany).

#### 6.2.3. Sectioning for LM and TEM

After embedding in Epon, we made a coarse trimming of the blocks using manually cut (7800 Knifemaker, LKB; Bromma, Sweden) glass knives (Ultra Glass Knife Strips #71014, EMS; Washington, PA, USA) on an ultramicrotome (Ultracut UCT or EM UC7; Leica Microsystems, Wetzlar, Germany). For LM, 1 µm-thick sections were cut with an 8.0 mm diamond knife (Diatome Histo AT 110, 8.0 mm; Nidau, Switzerland), mounted on microscopy slides (SuperFrost®Plus, 25x75x1 mm; ThermoScientific, Menzel-Gläser, Braunschweig, Germany), and set to dry at 50 °C on a heatbed (Medite OTS 40; Burgdorf, Germany). The mounted slides were then exposed to specific histological staining of myelin sheets with p-phenylenediamine (PPD; 1% w/v in a 50:50 solution of methanol:isopropanol) for 3 min, and briefly washed in acetone.

To measure axon diameters and myelin from TEM, we obtained ultra-thin sections (~ 40-70 nm) from the Epon-embedded samples using an ultramicrotome (Ultracut UCT or EM UC7, Leica Microsystems, Wetzlar, Germany). The sections were mounted on copper slot grids with a FORMVAR membrane. To obtain proper contrast to myelin for TEM, the sections are automatically stained on the grids with uranyl acetate and lead citrate (EM AC20; Leica Microsystems, Wetzlar, Germany).

### *6.3. Microscopy and image segmentation*

#### 6.3.1. LM of Epon-embedded samples

We initially obtained overview images (2752 x 2192 px, Lumenera; MBF Bioscience, Williston, VT, USA) of the Ø = 1 mm samples on LM (Olympus BX 60; Olympus Europe, Hamburg, DE) with a 10x lens (NA: 0.30; UPlanFI; Olympus).

#### 6.3.2. TEM of Epon-embedded samples

TEM images were obtained on a CM100 microscope (Philips; FEI, Eindhoven, NL). The CC sample, approximately encompassing the M1 axon tracts, was imaged at 4200x magnification in a sequential manner to cover the entire CC cross section in the Ø=1mm punctured sample. This resulted in images (2048 x 2048 pixels per image; 12 nm/pixel; ~24.6 x 24.6 µm) numbering from 16 from the top part of CC (N=1) to 69-77 images (N=4), which were acquired in an unbiased manner throughout the sample. We took care to avoid overlap of the images, any such regions from the first animal (N=1) were manually discarded from the stitched super image ("Stitching Tool" [23] in FIJI) to avoid overcounting of the same axons. The segmentation of the acquired images was obtained using AxonDeepSeg [24], using the built-in pre-trained model for TEM images. The model provided individual inner (d) axon diameters based on the segmented area and assuming the circular cross sections, as well as the corresponding g-ratios of all segmented axons. The acquired axon diameter distributions were filtered between 0.25 and 5.0 μm, based on visual inspection.

#### 6.3.3. TEM of Cryo-fixed tissue samples

TEM images of the cryo-fixed and Epon-embedded samples were obtained using the same microscope as above (CM100, Philips; FEI, Eindhoven, NL), with a pixel size of 8.55 nm and field of view (FOV) spanning 2048 x 2048 pixels, thus corresponding to approximately 17.5 x 17.5 μm. Fifteen TEM images were obtained from different positions within the same section and were then aligned relative to each other on a lower magnification image. The automatic segmentation method did not immediately work for cryo-TEM images due to their differing image contrast. Therefore, we manually segmented 607 axons in ITK Snap (RRID:SCR_002010) using the polygon tool and the adaptive paintbrush tool. The intra-axonal space was here defined as the area bounded by the innermost myelin layer. A connected components analysis of the segmented axons was then performed in MATLAB 2020a. Owing to the eccentricity of many of the axonal cross sections, the minor axis of an ellipse was used to represent axon diameter. The obtained axon diameter distributions were filtered between 0.1 μm and 5.0 μm, resulting in 603 axons. This lower diameter detection was feasible for the highly detailed cryo-TEM images, which allowed measurement of myelinated axons below 0.25 μm diameter in contrast to the Epon-TEM images

### 7. Calculating the structure-function relationship between conduction velocity and axon diameter

From the obtained length of the projection pathway (L) and the measured TCT, we calculated the estimated conduction velocity (CV) as follows:

$$\mathbf{CV = L / TCT} \qquad \mathbf{(1)}$$

Similarly, we calculated the conduction velocity along the pathways from the measured g-ratio and axon diameter, given Hursh's structure-function relationship [25–27]:

$$\mathbf{CV = (5.5/g) * d} \qquad \mathbf{(2)}$$

where g, i.e., the g-ratio, was set to group mean of EM data results, rather than the literature value of ~0.7 [14,27,28]. Hence, combining Eq. (1) and (2) allows us to predict the TCT from structural measures, or the converse prediction of axon diameter from the functional measure.

### 8. Statistical Analysis

#### *8.1. Opto-ElPhys*

Signal data from the two channels of the stereotrodes were averaged for each trial. For peak detection, all 150 trials for each condition were averaged. The coefficient of variation was calculated as the standard deviation (SD) of the signal amplitude of all individual trials at the exact sample of the peak latency, divided by the mean signal amplitude at the peak latency. A condition was considered reliable when the coefficient of variation of the trials was less than 1.00. Median latencies of all robust conditions were used to represent the latency of each animal, presented in violin plots (Fig. 1b). Medians of all animals along with whisker plots of most extreme values are indicated with vertical lines.

#### *8.2. Axon diameter distributions from histology*

To quantify the axon diameter distributions of the measured diameters from histology, we fitted gamma distributions with parameters *a* (shape), and *b* (scale), and used a location parameter of 0. Statistical features were expressed as the mean, SD, and the mode. The voxel-wise dMRI diameter estimates are weighted towards larger diameters, in part due to their larger volumes [19,29] and in part due to the nature of the scaling of the signal attenuation with diameter [12,30,31]. As such, the arithmetic means of the histological axon diameter distribution will not match the diameter index measured with dMRI. We accounted for this weighting by calculating the weighted diameter ($d_w$) from the histological axon diameter distributions. The weighting depends on the dMRI protocol used and upon the axon diameter distribution. In the wide pulse limit, $\delta \gg R^2/D_0$ [12,30,31], where $R$ is the axon radius from histology, $D_0$ is the intrinsic diffusivity and $\delta$ is the pulse-width of the diffusion encoding. In practice, this wide-pulse limit applies to most axons in the rat brain [8] Using this assumption we can calculate $d_w$ as [12,30]:

$$\mathbf{d_w = 2\left(\frac{<R^6>}{<R^2>}\right)^{0.25}} \quad \textbf{(3)}$$

**Supplementary Figures**

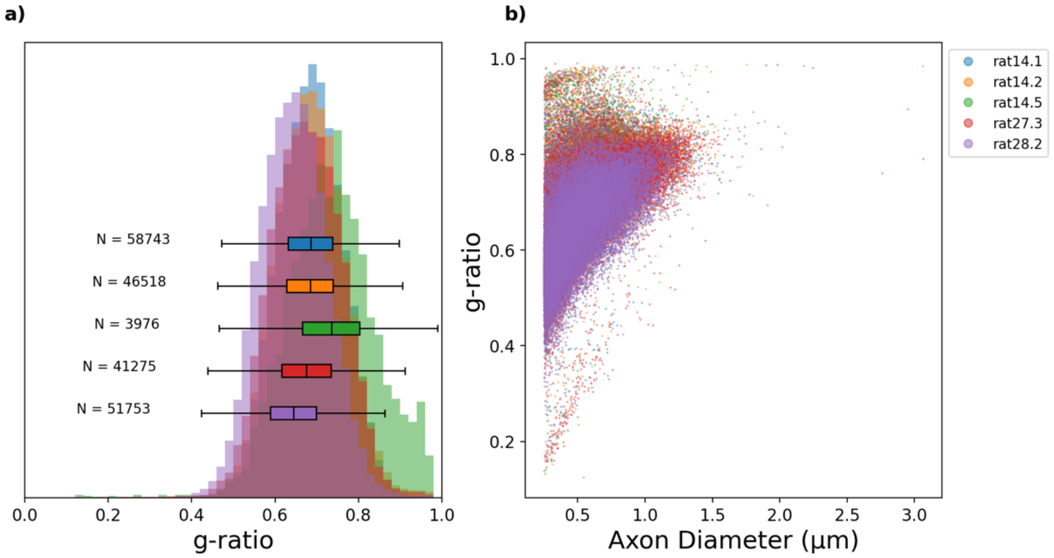


**Fig. S1.** Suppl. Figure 1: Axonal g-ratios, segmented from Epon-TEM. Distribution of g-ratios of the automatically seg-mented axons (Left panel). G-ratio vs axon diameter (Right panel).

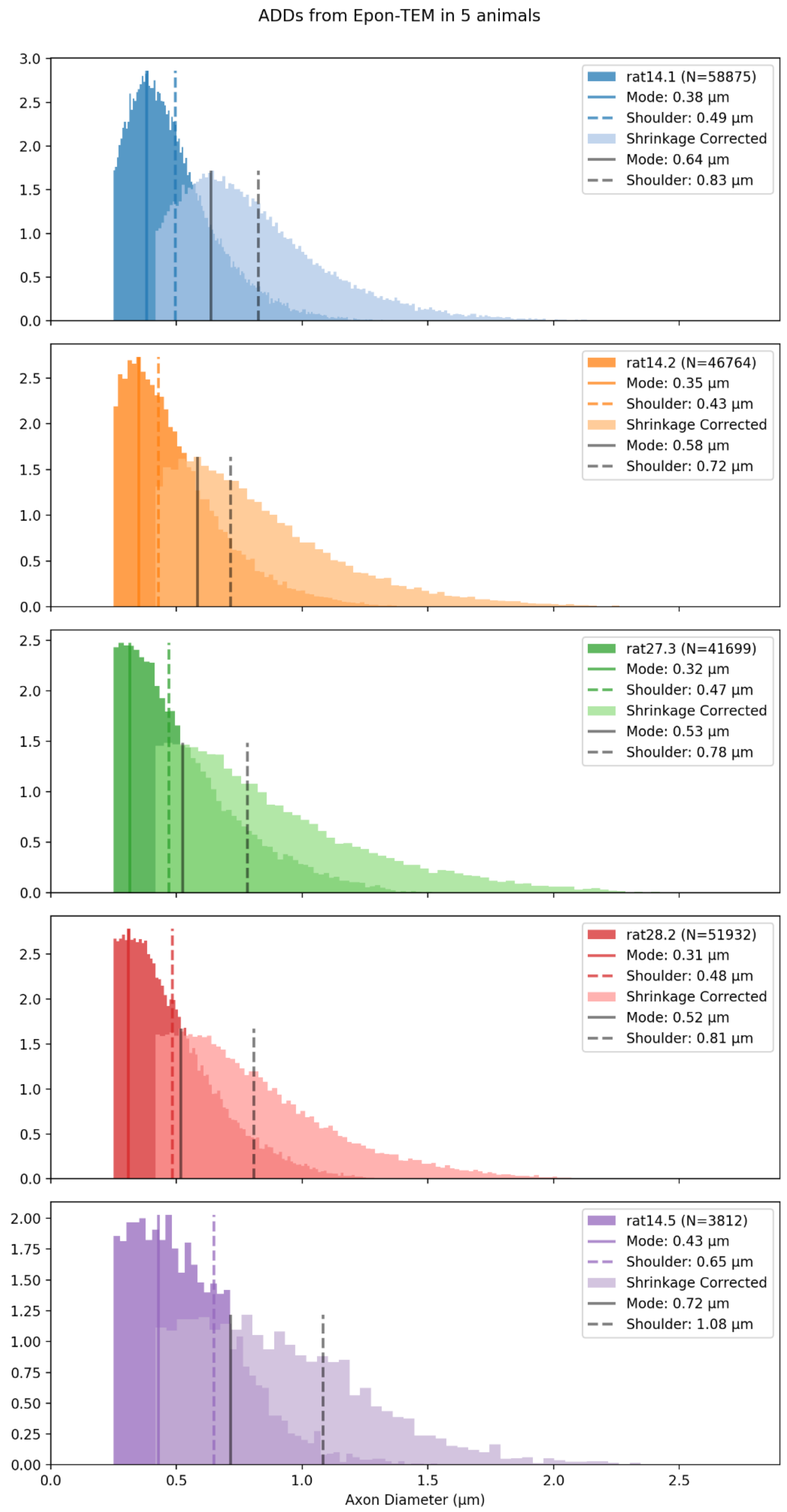


**Fig. S2.** a) Overview of axon diameters from segmented TEM images from EPON-embedded CC-tissue of individual rats. Mode and shoulder values were obtained visually.

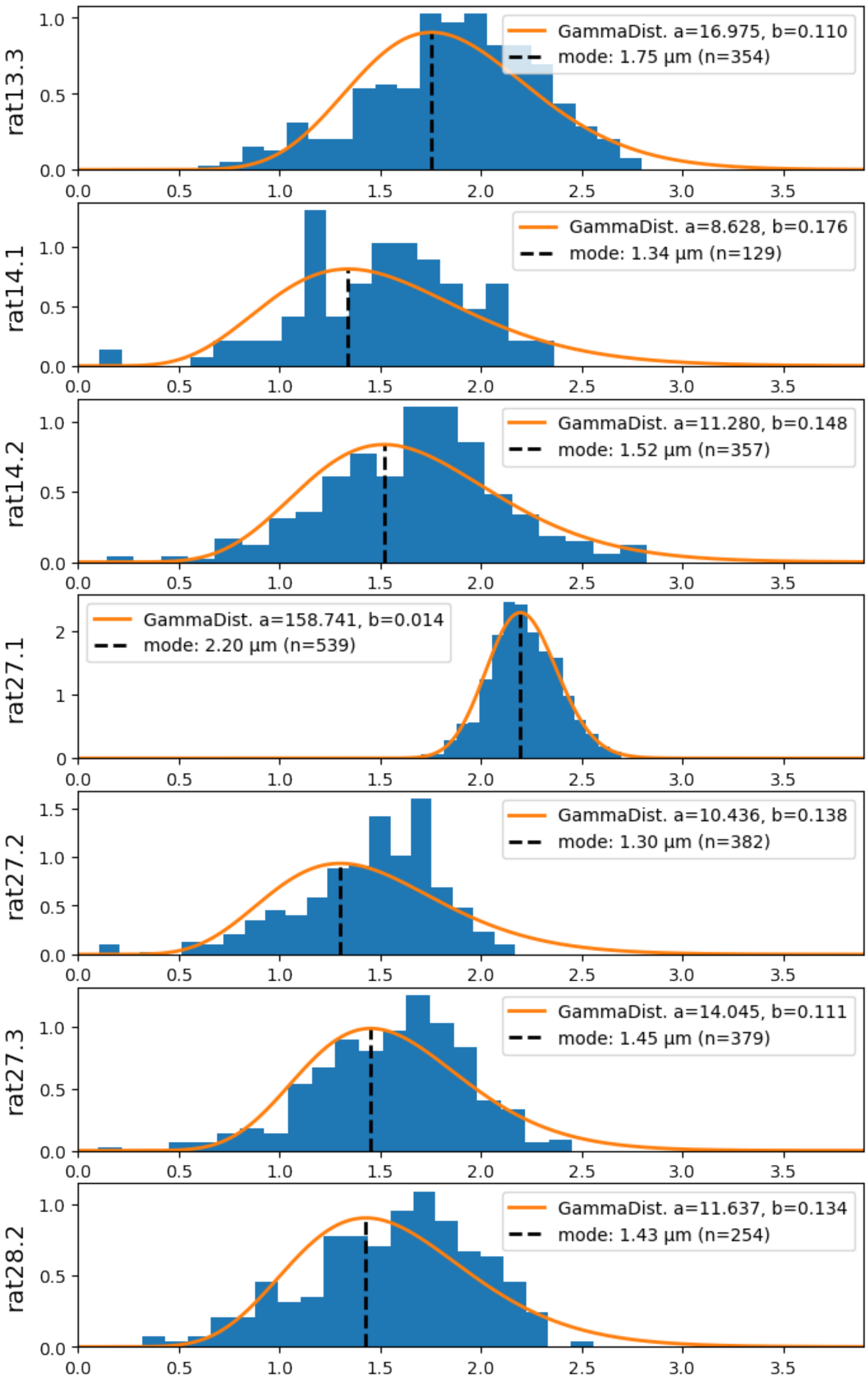


**Fig. S3:** Overview of predictions of axon diameters from dw-MRI of individual rats. N=number of extracted voxels, defined by the tractography streamlines projecting between the bilateral M1s and passing through the midsagittal plane.

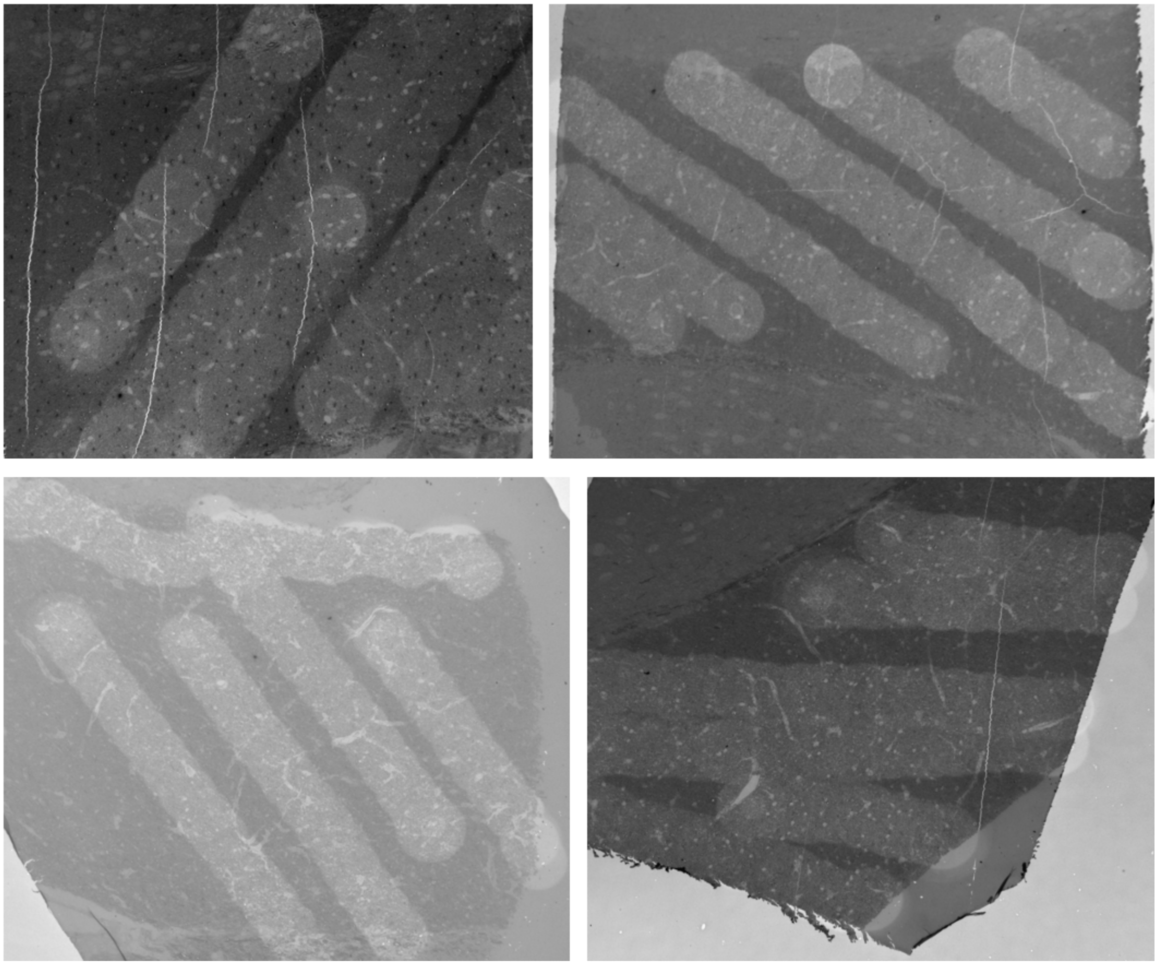

**Fig. S4:** TEM overview images at low magnification of the acquisition traverse paths at high magnification (4200x) from four rats. The overall samples correspond to ultrathin sections of the Ø=1mm punctures. The darker band in the middle of the sample corresponds to the corpus callosum. Each bleached circle corresponds to a focus area for image acquisition. The focus bleached circles are larger than the actual square field of view (2048x2048 pixels) and care was taken to avoid overlap of the images. The images were acquired in an unbiased manner by an other-wise uninvolved TEM-professional.

**Table S1.** Latencies from P1 and N1 for all animals

| Animal | P1 Peak | | | N1 Onset | | | N1 Peak | | |
|---|---|---|---|---|---|---|---|---|---|
| | Median (ms) | SD (ms) | Vals (N) | Median (ms) | SD (ms) | Vals (N) | Median (ms) | SD (ms) | Vals (N) |
| rat27.1 | 4.83 | 0.10 | 2 | 6.03 | 0.42 | 9 | 10.27 | 0.44 | 9 |
| rat27.2 | 5.87 | 0.27 | 22 | 7.63 | 0.36 | 24 | 11.90 | 0.50 | 24 |
| rat27.3 | 4.30 | 0.12 | 19 | 5.57 | 0.18 | 23 | 9.30 | 0.30 | 23 |
| rat28.2 | 7.07 | 0.42 | 12 | 8.94 | 0.44 | 12 | 13.62 | 0.62 | 12 |
| rat33.1 | - | - | 0 | - | - | 0 | - | - | 0 |
| rat35.2 | 3.30 | 0.21 | 3 | 4.73 | 0.35 | 15 | 10.93 | 0.53 | 15 |
| rat35.3 | 6.03 | 0.39 | 11 | 8.83 | 0.44 | 11 | 12.70 | 0.62 | 11 |
| rat35.4 | 7.42 | 0.08 | 2 | 10.52 | 0.15 | 2 | 15.68 | 0.35 | 2 |
| rat9.4 | - | - | 0 | - | - | 0 | - | - | 0 |
| rat10.1 | 5.40 | 0.22 | 7 | 6.60 | 0.29 | 8 | 10.79 | 0.76 | 8 |
| rat13.2 | - | - | 0 | 5.05 | 0.34 | 10 | 9.62 | 0.37 | 10 |
| rat13.3 | 7.32 | 0.34 | 4 | 8.87 | 0.28 | 4 | 14.18 | 0.77 | 4 |
| rat14.1 | 3.40 | 0.26 | 3 | 5.03 | 0.51 | 15 | 9.93 | 0.49 | 15 |
| rat14.2 | 3.57 | 0.25 | 4 | 5.87 | 0.28 | 22 | 10.45 | 0.43 | 22 |
| rat14.3 | 6.87 | 0.39 | 16 | 8.90 | 0.36 | 16 | 12.98 | 0.75 | 16 |
| rat14.5 | 6.98 | 0.32 | 14 | 8.58 | 0.28 | 14 | 12.87 | 0.66 | 14 |
| **median** | **5.87** | | **13** | **7.12** | | **14** | **11.42** | | **14** |
| **Mean ± Mean SD (within)** | **5.57** | **0.26** | **13** | **7.22** | **0.33** | **14** | **11.80** | **0.54** | **14** |
| **Mean ± SEM** | **5.57** | **0.43** | **13** | **7.22** | **0.50** | **14** | **11.80** | **0.51** | **14** |
| **Mean + SD (Between)** | **5.57** | **1.49** | **13** | **7.22** | **1.81** | **14** | **11.80** | **1.84** | **14** |

**Table S2.** Electrode and fiber depths measured from T2 FISP 3D MRI

| RatID | ScanID | Electrode Depth (left) (mm) | Fiber Depth (right) (mm) |
|---|---|---|---|
| rat33.1 | M0613 | -0.97 | -0.97 |
| rat13.2 | M0642 | -1.10 | -1.22 |
| rat13.3 | M0649 | -1.01 | -1.33 |
| rat14.1 | M0620 | -1.07 | -1.23 |
| rat14.2 | M0641 | -1.07 | -0.94 |

**Table S3.** Transcallosal pathway length between bilateral M1s, from tractography of dw-MRI

| RatID | ScanID | Left Tract (mm) | Right Tract (mm) | Adjust H (mm) | Adjust mid CC (mm) | Total (mm) |
|---|---|---|---|---|---|---|
| rat13.2 | M0642 | 7.58±2.40 | 5.70±2.23 | 0 | 0.23 | 13.51 |
| rat13.3 | M0751 | 3.52±0.76 | 6.54±1.81 | 1.012 | 0.23 | 11.30 |
| rat14.1 | M0765 | 5.64±1.53 | 6.46±1.87 | 0 | 0.23 | 12.33 |
| rat14.2 | M0773 | 5.50±1.26 | 5.78±1.29 | 0 | 0.23 | 11.51 |
| rat33.1 | M0613 | 4.98±1.16 | 4.45±0.80 | 0 | 0.23 | 9.66 |
| rat27.2 | M0781 | 5.39±0.73 | 4.61±0.67 | 0 | 0.23 | 10.23 |
| rat27.3 | M0783 | 5.07±0.83 | 5.15±0.63 | 0 | 0.23 | 10.45 |
| rat28.2 | M0787 | 5.74±1.59 | 6.83±2.23 | 0 | 0.23 | 12.80 |
| **AVG** | | | | | | **11.47 ± 0.47** |

**Table S4.** Non-exhaustive overview of histological investigations of axon diameter distribution in CC of Rats.

| Sex | Age | Region | Value (µm) | Metric | Imaging | Tissue state | Study |
|---|---|---|---|---|---|---|---|
| M | P150 | Anterior CC<br>Middle CC<br>Poterior | ~0.58<br>~0.56<br>~0.50 | mean | TEM<br>16,500x | Perf.F: 4% FA + 1% Glut, Dehyd., Resin (should you identify abbreviations in table caption?) | [32] |
| M | P58 | Forceps Minor | ~0.89 | mean | LM<br>100x (oil) | Perf.F: 1% FA + 1.25% Glut<br>Dehyd., Resin | [33] |
| M | P120 | #1: Genu<br>#2<br>#3: Midbody<br>#4<br>#5: Splenium | ~0.9<br>~0.6<br>~1.0<br>~1.1<br>~0.6 | mode<br>(read from axon diameter distributions) | TEM<br>15,000x | Perf.F: FA+Glut<br>PostF: 2.5% Glut,<br>Dehyd., Resin | [34] |
| M + F | P70 | Splenium | ~0.5 | mean | TEM<br>20,000x | Perf.F: 1% FA + 1.0% Glut<br>PostF: 2% FA + 0.1% Glut<br>Dehyd., Resin | [35] |
| F | P84 | Genu | ~1.22 | median | Confocal LM, 63x | Perf.F: 4% FA<br>PostF: 4% FA<br>Hydrated | [12] |
| M+F | Young adult | Splenium | [0.11 - 0.2] | modal bin<br>(binned interval) | TEM<br>33,000x | Perf.F: 4% Glut<br>PostF: 4% Glut<br>Dehyd., Resin | [36] |

14. Andersson M, Kjer HM, Rafael-Patino J, Pacureanu A, Pakkenberg B, Thiran J-P, et al. Axon morphology is modulated by the local environment and impacts the noninvasive

investigation of its structure-function relationship. Proc Natl Acad Sci U S A. 2020;117: 33649–33659. doi:10.1073/pnas.2012533117
